# Stabilized supralinear network dynamics account for stimulus-induced changes of noise variability in the cortex

**DOI:** 10.1101/094334

**Authors:** Guillaume Hennequin, Yashar Ahmadian, Daniel B. Rubin, Máté Lengyel, Kenneth D. Miller

## Abstract

Variability and correlations in cortical activity are ubiquitously modulated by stimuli. Correlated variability is quenched following stimulus onset across multiple cortical areas, suppressing low-frequency components of the LFP and of *V*_m_-LFP coherence. Modulation of Fano factors and correlations in area MT is tuned for stimulus direction. What circuit mechanisms underly these behaviors? We show that a simple model circuit, the stochastic Stabilized Supralinear Network (SSN), robustly explains these results. Stimuli modulate variability by modifying two forms of effective connectivity between activity patterns that characterize excitatory-inhibitory (E/I) circuits. Increases in the strength with which activity patterns inhibit themselves reduce correlated variability, while increases in feedforward connections between patterns (transforming E/I imbalance into balanced fluctuations) increase variability. These results suggest an operating regime of cortical dynamics that involves fast fluctuations and fast responses to stimulus changes, unlike previous models of variability suppression through suppression of chaos or networks with multiple attractors.

---

Neuronal activity throughout cerebral cortex is variable, both temporally during epochs of stationary dynamics and across repeated trials despite constant stimulus or task conditions (Softky and Koch, 1993; Churchland et al., 2010). Moreover, variability is *modulated* by a variety of factors, most notably by external sensory stimuli (Churchland et al., 2010; Kohn and Smith, 2005; Ponce-Alvarez et al., 2013), planning and execution of limb movements (Churchland et al., 2006, 2010), and attention (Cohen and Maunsell, 2009; Mitchell et al.,2009). Modulation of variability occurs at the level of single-neuron activity, e.g. membrane potentials or spike counts (Finn et al., 2007; Poulet and Petersen, 2008; Gentet et al., 2010; Churchland et al., 2010; Tan et al., 2014), but also in the patterns of joint activity across populations, as seen in multiunit activity or the local field potential (LFP) (Tan et al., 2014; Chen et al., 2014; Lin et al., 2015). Variability modulation shows stereotypical patterns: not only does the onset of a stimulus quench variability overall, and in particular correlated variability that is “shared” across many neurons (modeled as fluctuations in firing rates and typically found to be low-dimensional; Lin et al., 2015; Goris et al., 2014; Ecker et al., 2014, 2016; Churchland et al., 2010), but the degree of variability reduction can also depend on the tuning of individual cells. For example, in area MT, variability is quenched more strongly in cells that respond best to the stimulus, and correlations decrease more among neurons with similar stimulus preferences (Ponce-Alvarez et al., 2013; Lombardo et al., 2015). Although these patterned modulations of variability are increasingly included in quantitative analyses of neural recordings (Renart and Machens, 2014), it is still unclear what they imply about the dynamical regime in which the cortex operates.

Three different dynamical mechanisms have been proposed to explain some selected aspects of cortical variability. The so-called “balanced network” model (van Vreeswijk and Sompolinsky, 1998; Renart et al., 2010) has been highly successful at explaining in general the asynchronous and irregular nature of action potential firing in cortical neurons under normal operating conditions (Softky and Koch, 1993). However, very strong, very fast inhibitory feedback in the balanced network suppresses correlated rate fluctuations away from that stable state (van Vreeswijk and Sompolinsky, 1998; Renart et al., 2010;Tetzlaff et al., 2012), leaving only fast variability due to irregular spiking. Because the shared variability is already eliminated, stimuli cannot modulate that variability. This has been rectified in models in which not only the spiking of neurons but also their underlying firing rates are variable. In “attractor models”, the network noisily wanders among multiple possible stable states (“attractors”) in the absence of a stimulus, thus operating in a marginally stable state characterized by shared variability. Stimuli then suppress this shared variability by pinning fluctuations to the vicinity of one particular attractor (Blumenfeld et al., 2006; Litwin-Kumar and Doiron, 2012; Deco and Hugues, 2012; Ponce-Alvarez et al., 2013; Doiron and Litwin-Kumar, 2014; Mochol et al., 2015). In chaotic network models (Sompolinsky et al., 1988), strong firing rate fluctuations are typically low-dimensional (hence “shared”), and certain types of stimuli can suppress chaos, thus quenching across-trial variability (Molgedey et al., 1992; Bertschinger and Natschlger, 2004; Sussillo and Abbott, 2009; Rajan et al., 2010). While both the attractor and the chaotic mechanisms can explain the general phenomenon of stimulus-induced reduction of variability, only the former has been proposed to explain the stimulus-tuning of variability reduction – and even that required considerable fine tuning of parameters to keep it in the “metastable” regime, in which the system stays near attractors yet noise can move the system between them (Ponce-Alvarez et al., 2013).

Here we explored a qualitatively different model of cortical dynamics, the stabilized supralinear network (SSN; Ahmadian et al., 2013; Rubin et al., 2015). In the SSN, single neurons have supralinear input/output (I/O) curves (Priebe and Ferster, 2008), which yields a transition between two regimes at the level of circuit dynamics. For weak external inputs, network dynamics are stable even without inhibition. For stronger inputs, firing rates grow towards steeper parts of the I/O curves, leading to potential instability due to growing recurrent excitation, but feedback inhibition dynamically cancels the destabilising effect of this supralinearity, thus keeping the network in a fundamentally stable (as opposed to a metastable or chaotic) operating regime. This stabilization is achieved by a “loose” cancellation of moderately large E and I inputs, in contrast to the balanced network model, in which there is a precise cancellation of very large E/I inputs. We showed (Rubin et al., 2015) that the SSN naturally explains many cortical nonlinear behaviors, including sublinear summation of responses to different stimuli (“normalization”, Carandini and Heeger, 2012), surround suppression, and their nonlinear changes in behavior with stimulus strength. These behaviors cannot arise in the balanced network, because in that regime responses must be linear functions of the external input (though see Mongillo et al., 2012).Importantly, the SSN also presents a promising candidate for understanding variability modulation: its loose E/I balance is such that inhibitory feedback is weak enough for shared network variability to subsist over a broad range of input strengths, and we also expect its nonlinear collective behaviour to lead to a non-trivial modulation of this shared variability with the stimulus.

Here we show that, indeed, the SSN in the inhibition-stabilized regime increasingly and gradually suppresses correlated rate variability with increasing external input strength, rather than eliminating it like the balanced network. As a result, the SSN naturally and robustly explains modulation of cortical variability, including its tuning dependence. We first analyzed variability in the simplest stochastic instantiation of the SSN, with two unstructured populations of excitatory (E) and inhibitory (I) cells, and found that an external stimulus could strongly modulate the variability of population activities. In particular, the model predicts stimulus-induced quenching of variability, as well as a reduction of the low-temporal-frequency coherence between local population activity and single-cell responses, as found experimentally (Poulet and Petersen, 2008; Churchland et al., 2010; Chen et al., 2014; Tan et al., 2014). Furthermore, tuning-dependent modulations of Fano factors and noise correlations by stimuli arise robustly in a more detailed architecture with structured connectivity, and are consistent with those found in area MT of the awake monkey (Ponce-Alvarez et al., 2013).

Mechanistically, input-induced modulation of variability in the SSN originates from input-dependent changes in effective connectivity between neurons, which themselves arise from the presence of nonlinear neuronal input/output functions. To dissect these mechanisms, we first analyzed a simple model of one E and one I population. We decomposed the effective connectivity into two types of input-dependent interactions between a pair of E and I activity patterns. One is a self-connection of an activity pattern onto itself, which more strongly suppresses variability as it becomes increasingly inhibitory, summarizing the effect of growing feedback inhibition. In the inhibition-stabilized regime, these connections grow more strongly inhibitory with increasing external input due both to the overall strengthening of effective connections and to the relatively faster growth of I vs. E firing rates, and thus of I vs. E effective connection strengths, that arises from the dynamics that keep the network stable. The other type of interaction is a feedforward connection from one activity pattern to another, which causes small differences between E and I cell activity to drive joint activity of E and I cells (“balanced amplification”, Murphy and Miller, 2009). These feedforward connections also grow with increasing external input strength, enhancing variability. We show that variability enhancements dominate for low levels of input (which might be below the levels of spontaneous activity), but suppression of variability via inhibitory feedback always dominates for larger inputs. The same insights generalized to a more complex architecture to explain the tuning-dependent reduction of variability by interactions between a small number of E and I activity pattern-pairs.

Our results have important implications beyond offering a new mechanistic understanding of cortical variability: the SSN is distinguished by dynamics in which the network responds to input changes on fast time scales comparable to those of isolated cells, rather than on much longer time scales created by recurrent excitation, which tend to govern dynamics in multi-attractor and chaotic networks (Murphy and Miller, 2009). This regime of fast fluctuations offers distinct computational advantages (Hennequin et al., 2014a), and seems to characterize at least mouse V1 (Reinhold et al., 2015).

## Results

Single neurons in sensory areas respond supralinearly to their inputs. Plotting momentary firing rates, *r*, versus average membrane potentials, *V*_m_, often reveals an approximate threshold power-law relationship (Figure 1B):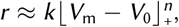, where *k* is some scaling constant; *V*_0_ ≈ −70 mV is a threshold that often approximates, and that we will always take equal to, the cell’s resting potential *V*_rest_; 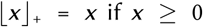otherwise; and the exponent *n* ranges from 1 to 5 in V1 (Priebe and Ferster, 2008). Importantly, this approximation is accurate over the entire dynamic range of neurons under normal spontaneous or stimulus-evoked conditions, i.e. neuronal responses rarely saturate at high firing rates. Accordingly, we modeled *V*_m_ dynamics as a simple low-pass filtering of synaptic inputs obtained as a weighted sum of presynaptic firing rates and external inputs (Experimental Procedures and SI):

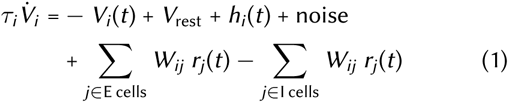

where *V*_*i*_ denotes the *V*_m_ of neuron *i*, *τ*_*i*_ is its membrane time constant (20 ms and 10 ms for excitatory and inhibitory cells, respectively), *V*_rest_ = −70 mV is a resting potential, *W*_*ij*_ is the (positive or zero) strength of the synaptic connection from neuron *j* to neuron *h_i_*(*t*) is the potentially time-varying but deterministic component of external input, and the momentary firing rate of cell *i* is given by

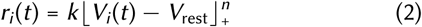

with *n* = 2 (Figure 1B; see also SI for an extension to other exponents). This is the stabilized supralinear network model studied in (Ahmadian et al., 2013; Rubin et al., 2015), but formulated with voltages rather than rates as the dynamical variables (the two formulations are mathematically equivalent when all neurons have the same time constant, Miller and Fumarola, 2011) and with noise added.

As experiments support Equation 2Equation 2 when both membrane potentials and spike counts are averaged in 30 ms time bins (Priebe and Ferster, 2008), *V*_m_ here stands for a coarsegrained (low-pass filtered) version of the raw somatic membrane potential, and in particular it does not incorporate the action potentials themselves. Thus the effective time resolution of our model was around 30 ms which allowed studying the effects of inputs that did not change significantly on timescales shorter than that. Accordingly, in Equation 1 we assumed that external noise had a time constant *τ*_noise_ = 50 ms, in line with membrane potential and spike count autocorrelation timescales found across the cortex (Azouz and Gray, 1999; Berkes et al., 2011; Murray et al., 2014).

We focused on analysing how the intrinsic dynamics of the network shaped external noise to give rise to stimulus-dependent patterns of response variability. We studied a progression of connectivity architectures **W** of increasing complexity, all involving two separate populations of excitatory and inhibitory neurons. We also validated our results in large scale simulations of spiking neuronal networks.

### Variability of population activity: modulation by external input

We first considered a simple circuit motif: an excitatory (E) unit and an inhibitory (I) unit, recurrently coupled and receiving the same mean external input *h* as well as their own independent noise (Figure 1A). In this simple network, the two units represent two randomly connected populations of E and I neurons, a canonical model of cortical networks (Vogels et al., 2005). Thus, their time-varying activity, *V*_E_(*t*) and *V*_i_(*t*), represent the momentary population-average membrane potential of all the E and I cells respectively. While these population-level quantities cannot be compared directly with the intracellularly recorded membrane potentials of individual cells, we used their average to model the extracellularly recorded LFP. Despite its simplicity, this architecture accounted well for the overall population response properties in the larger networks with more detailed connectivity patterns that we analyzed later.

**Figure 1.**
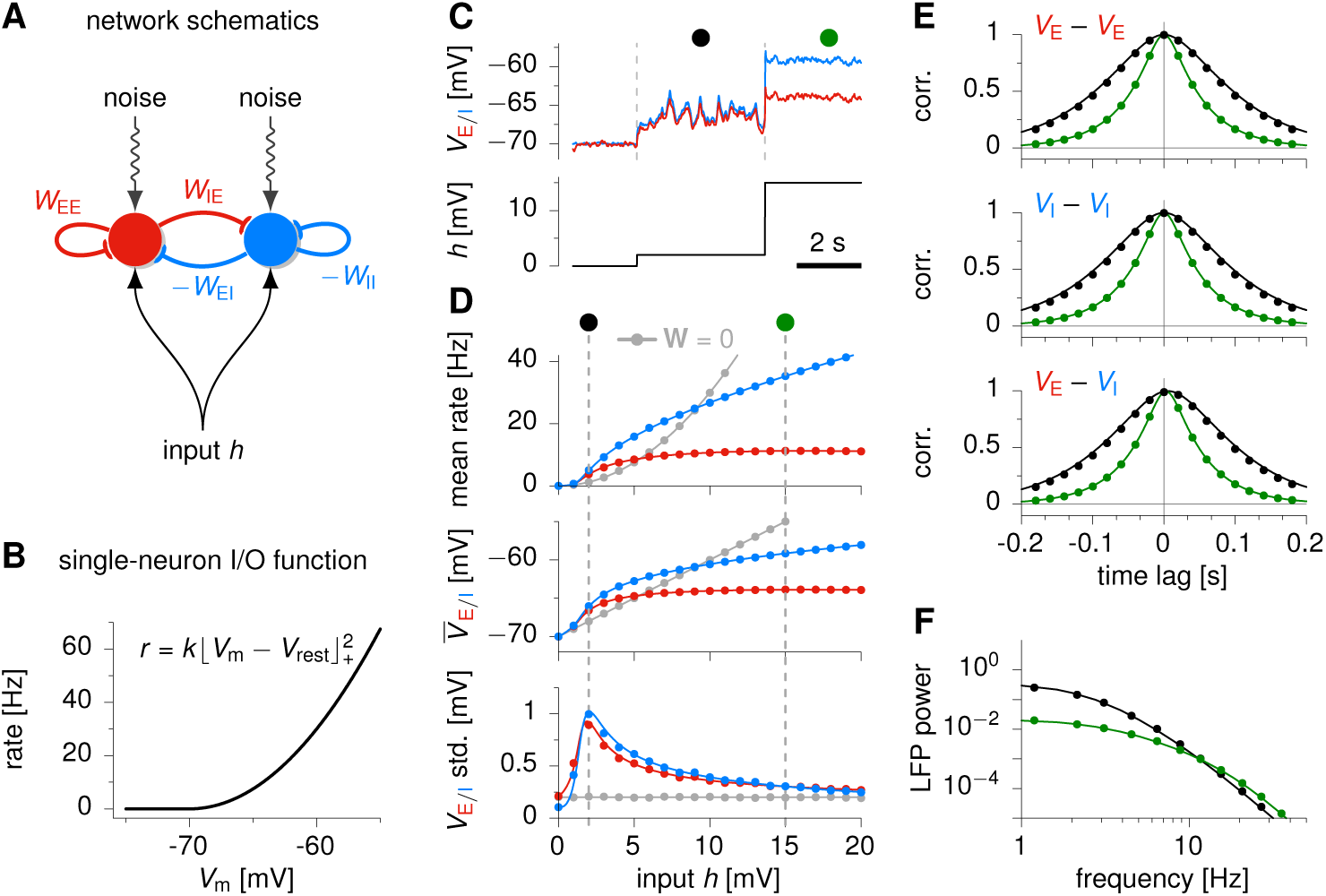
Activity variability in a reduced, two-population model of a supralinear stabilized network. (**A**) The network is composed of two recurrently connected units, summarizing the activity of two populations of excitatory (red) and inhibitory (blue) neurons. Both units receive private noise and a common constant input *h*. (**B**) Threshold-quadratic gain function determining the relationship between membrane potential and momentary firing rate of model neurons (Equation 2).(**C**) Sample *V*_E/I_ traces for both units (top), as the input is increased in steps from *h* = 0 to 2 mV to 15 mV (bottom). (**D**) Dependence of population activity statistics on stimulus strength *h*. Top: mean E and I firing rates; middle: mean *V*_E/I;_ bottom: standard deviation of *V*_E/I_ fluctuations. The comparison with a purely feedforward network (**W** = 0) is shown in gray. (**E**) Population Vm auto- and cross-correlograms in stationary conditions, when *h* = 2 mV and *h* = 15 mV (black and green, respectively, cf. marks in panels C-D). In both input conditions, *V*_E_ and *V*_I_ fluctuations are highly correlated, inhibition lagging behind excitation by a few ms. Note also that *V*_E/I_ fluctuations are faster for *h* = 15 mV. (**F**) LFP power spectrum for low input (*h* = 2 mV) and high input (*h* = 15 mV) conditions. The LFP is modelled as an average of *V*_E_ and *V*_I_, weighted by assumed relative population sizes (80% E,20% I). Strong input mostly suppresses low frequencies. In (D), (E) and (F), dots show the results of 1000 second-long numerical simulations of Equation 1, and solid lines show theoretical predictions derived analytically using novel nonlinear techniques (Hennequin and Lengyel, *in prep*.)

The connectivity matrix in this reduced model takes the form

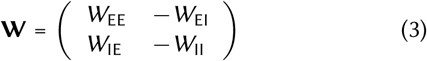

where *W*_*AB*_ is the magnitude of the connection from the unit of type *B* (E or I) to that of type *A*. The W terms were chosen such that the collective dynamics of the network remained stable for any input despite the strongly supralinear input-output functions of individual neurons (Equation 2, Figure 1B; see also Experimental Procedures).

Activity in the network exhibited temporal variability due to the noisy input. We found that the external, steady input *h* strongly modulated both the mean, *V̅*_E/I_, and the (co)variance of the fluctuations in *V*_E_ and *V*_I_ (Figure 1C-E). When *h* = 0, there was no input to drive the network, and *V*_E_ and *V*_i_ hovered around *V*_rest_ = −70 mV, fluctuating virtually independently with standard deviations essentially matching those that would arise without recurrent connections (**W** = 0, gray line in Figure 1D, bottom). For a somewhat larger input, *h* = 2 mV, both E and I populations fired at moderate rates (3-4 Hz) (Figure 1D,top), but now also exhibited large and synchronous population *V*_m_ fluctuations (Figure 1C, black circle mark). For yet larger inputs (*h* = 15 mV), fluctuations remained highly correlated but were strongly quenched in their magnitude (Figure 1C, green circle mark).

Figure 1D shows how the temporal (or, equivalently, the across-trial) mean and variability of activities varied over a broad range of input strengths. We observed that, with growing external input, population mean *V*_m_ grew linearly or supralinearly for small inputs, but for larger inputs grew strongly sublinearly, with *V̅*_I_ growing faster than *V̅*_E_ (Figure 1D, middle; Ahmadian et al., 2013; Rubin et al., 2015). Variability in both *V*_E_ and *V*_i_ typically increased for small inputs, peaking around this transition between supralinear and sublinear growth, and then decreased with increasing input (Figure 1D, bottom). These effects were robust over a broad range of network parameters (gain functions, connection weights, input gains and correlations), as long as they ensured dynamical stability (Supplementary Figures S1 and S2). Although the precise amplitude and position of the peak of *V*_m_ variance depended on network parameters, the overall non-monotonic shape of variability modulation was largely conserved. In particular, we could show analytically that variability suppression occurs earlier (for smaller input *h*) in networks with strong connections, or, for fixed overall connection strength, in networks that are dominated by feedback inhibition (*W*_EI_*W*_IE_ ≫ *W*_EE_ *W*_II_; SI). More generally, we found that the firing rates at the peak of variability are typically low (2.5 Hz on average over a thousand randomly parameterized stable networks, and below 6 Hz for 90% of them; cf. SI). Since these rates are comparable to cortical spontaneous firing rates, this predicts that increased sensory drive should generally result in variability quenching in cortical LFPs.

Importantly, input-modulation of variability required recurrent network interactions. This was revealed by comparing our network to a purely feedforward circuit (**W** = 0) which exhibited qualitatively different behaviour (Figure 1D, gray). In the feedforward circuit, mean *V*_m_ remained linear in *h*, so that mean rates rose quadratically with *V*_m_ or *h*, and fluctuations in *V*_m_ no longer depended on the input strength.

### Changes in effective connectivity shape variability in the SSN

The effects of input *h* on variability could be understood from the way it modified the *effective connectivity* of the circuit. An effective connection quantifies the impact of a small momentary change in the *V*_m_ of the presynaptic neuron on the total input in its postsynaptic partner. Formally, we derived effective connectivity from a linearization of Equations 1 and 2: we start with the steady state mean voltage *V̅*_i_ for the given mean input *h*, and analyze the dynamics of each neuron’s small, momentary, noise-induced deviations *δV*_i_(*t*) from *V̅*_i_:

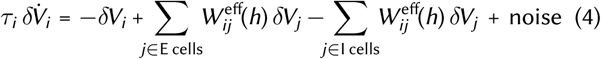

where the effective connection strength,

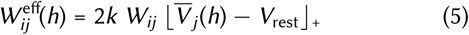

was proportional to the mean activation of unit *j,* which itself depended on the input *h* as seen above (cf. Figure 1D, middle). This growth of effective connectivity with increasing *V̅* arose because of the supralinear input/output function: the effective connectivity 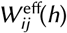 is the biophysical weight *W*_ij_ multiplied by the gain of the presynaptic cell – the change in its firing rate per change in its voltage – which is the ever-increasing slope of its input/output function (Figure 1 B).

How are changes in effective connectivity translated into changes in variability? For zero input, *h* = 0, the effective connections are zero, so we should expect behavior as if the neurons were uncoupled (**W** = 0), as observed (Figure 1D, compare blue and red lines with gray lines at *h* = 0). With increasing *h*, the effective connectivity strengthens, but – as growth of *V̅* becomes sublinear – grows more rapidly for inhibitory than for excitatory weights, reflecting the faster growth of *V*_I_ over *V*_E_ (Figure 1D, middle). This greater relative growth of inhibitory response is a robust outcome of the network maintaining stability despite increasing effective connectivity (Ahmadian et al., 2013; Rubin et al., 2015). These changes in effective connectivity can have conflicting effects: the increasingly strong weights can increase excitatory or driving effects that amplify fluctuations and increase variability (Murphy and Miller, 2009), but they and the relatively stronger inhibition also increase inhibitory effects, suppressing fluctuations and decreasing variability (Renart et al., 2010; Tetzlaff et al., 2012). The actual behavior of the network was mixed: variability first increased and then decreased as the input grew (Figure 1D, bottom). This suggests a changing balance of the variability-amplifying and -attenuating effects of changing effective connectivity.

What determines this changing balance? To study this, we examined the flow of *V*_m_ trajectories, visualized in the plane of joint *δV*_E_ and *δV*_I_ fluctuations (Figure 2A-C). In general, *V*_m_ trajectories underwent diffusion driven by the external input noise (Figure 2A, gray trajectory). With no external mean drive (*h* = 0), effective connectivity being negligible, the only contribution to the flow of activity was the leak in the E and I populations – the − *δV_i_* term in Equation 4 (Figure 2A, green arrows). Leak created a restoring “force field” by pulling both E and I activities back towards rest with their characteristic time constants (Figure 2A, green arrows, growing linearly as one moves away from the origin against their pointing direction) and thus contained diffusion so that it had a finite (co)variance (Figure 2A, black ellipse).

**Figure 2.**
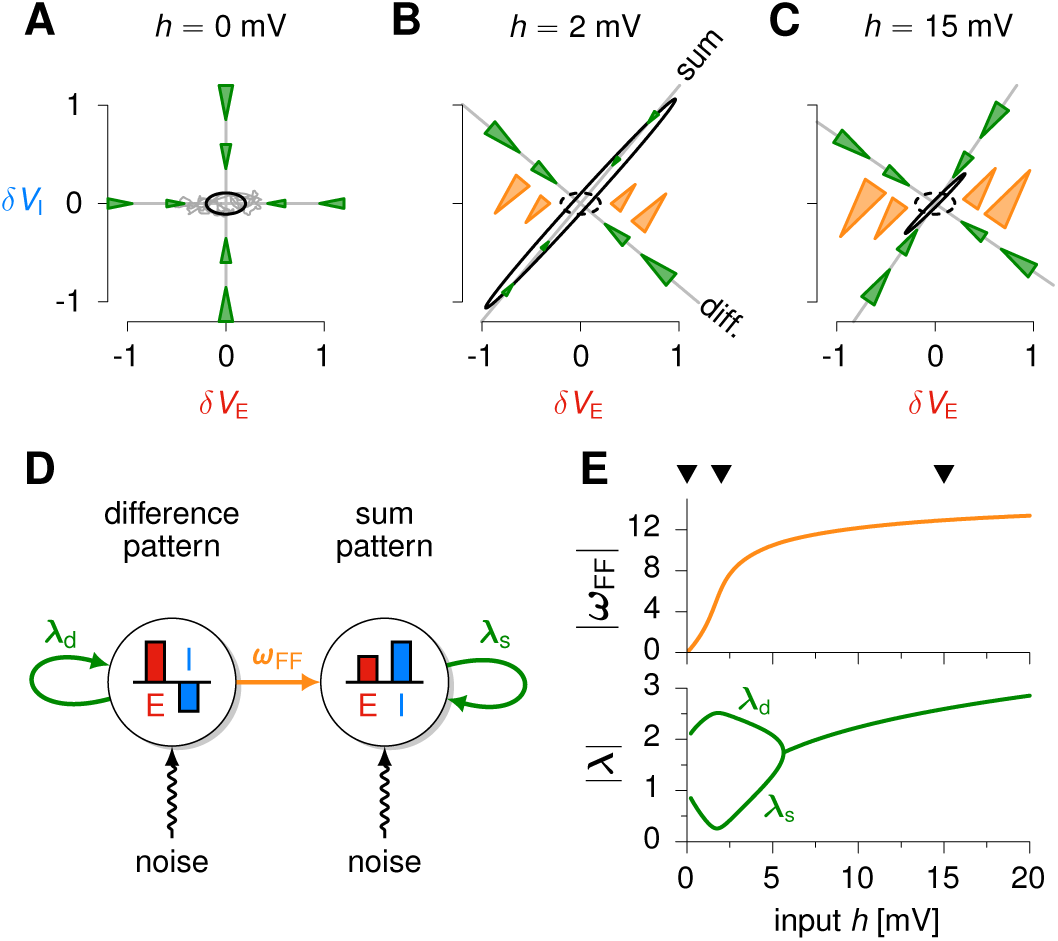
The origins of input-dependent modulation of variability. **(A-C)** Visualization of the influence of single-neuron leak and effective connectivity on the (co-)variability of E/I activity in the two-population SSN of Figure 1. In (**A**), *h* = 0, so the only contributor to the flow of trajectories is the leak in each population (green force field acting along the cardinal axes of E/I fluctuations – the flow is more compressive along the I axis due to the shorter membrane time constant in I cells). This flow contains the diffusion due to input noise (cf.example trajectory in gray), resulting in uncorrelated baseline E/I fluctuations (black ellipse – contour line of the joint normal distribution of *δV*_E_ and *δV*_I_ at one standard deviation). In (**B-C**), the network is driven by a non-zero *h*, and the effective recurrent connectivity adds to the leak to instate two types of force fields steering fluctuations: a restoring force field (green, generalizing the leak in (A)) and a shear force field (orange). The relative contributions of the two force fields determine the size and elongation of the E/I covariance (solid black ellipse – contour line of the joint normal distribution of *SV*_E_ and *SV*_I_ at one standard deviation). In (**B–C**)the network is driven by a non-zero *h*, and the effective recurrent connectivity adds to the leak to instate two types of force fields steering fluctuations: a restoring force field (green, generalizing the leak in (A)) and a shear force field (orange). The relative contributions of the two force fields determine the size and elongation of the E/I covariance (solid black ellipses). The black ellipse in (A) is reproduced in (B–C) for comparison (dashed ellipses). Triangular arrows are proportional in area to the contribution they make to the total flow of fluctuations. The origin (δV = 0) corresponds to stationary mean population activity for the given input strength h (see labels). (**D**) Illustration of the decomposition of the effective connectivity (for a given mean stimulus *h*) as couplings between a difference-like pattern (left) and a sum-like pattern (right; cf. rotated gray axes in (B-C)). For a given input *h*, the difference feeds the sum with weight *ω*_FF_ (orange arrow), and the difference and sum patterns inhibit themselves with negative weight λ_d_ and λ_s_ respectively (green arrows). These *h*-dependent couplings scale the corresponding force fields in (A-C) (note color consistency). (**E**) Input-dependence of |ω_FF_| (top, orange) and |λ_d_| and |λ_s_| (bottom, green).

With increasing mean external drive *h*, the effective connectivity of the network also began to contribute to the dynamics and thus to the total flow. While the connectivity between E and I populations was fully recurrent, it could be conveniently decomposed into a set of simpler interactions among a pair of joint E-I activity patterns, one a weighted difference and the other a weighted sum of E and I activities (rotated gray axes in Figure 2B-C; Murphy and Miller, 2009, see also Supplementary Figure S2E). First, both patterns inhibited themselves through negative self-couplings λ_d_ and λ_s_ (Figure 2D, green arrows). These “restoring forces” included the effects of both leak and recurrent feedback, and acted along the sum and difference axes now, rather than on E and I cells separately (compare green arrows between Figure 2A and B). Second, the difference pattern fed the sum with an effective feed-forward coupling *ω*_FF_ (Figure 2D, orange arrow). This effect, known as balanced amplification (Murphy and Miller, 2009; Hennequin et al., 2014b), created a “shear” force field (Figure 2B-C, orange arrows, growing linearly along the difference axis, but unchanged by movement along the sum axis) acting on *V*_m_ fluctuations such that excursions away from E-I balance (movements along the difference axis) were transported along the sum direction.

While the purely restorative force field at *h* = 0 shaped network variability simply by containing diffusion (Figure 2A), the combination of shear and restoring forces at *h* > 0 steered diffusion differentially along the sum and difference directions, resulting in various patterns of correlated E/I *V*_m_ variability (Figure 2B-C, black ellipses). Importantly, these forces depended on the input (compare Figure 2B and C) as their magnitude was scaled by the coupling terms characterizing effective connectivity, *ω*_FF_, λ_d_ and λ_s_, which in turn fundamentally depended on the input (Equation 5, Figure 2E). This is the origin of input-dependent variability in the SSN.

In the small-input regime, we found that the feedforward coupling *ω*_FF_ typically grew quickly (Figure 2E, orange) whereas the negative self-couplings λ_s_ and λ_d_ tended to grow more slowly, or to even weaken transiently (Figure 2E, green; this transient weakening of self-coupling was atypical in randomly instantiated networks, SI). These two effects combined to yield an initial increase in *V*_m_ variability for increasing external drive. For example, for the particular parameters used in the simulations shown in Figures 1 and 2, there was little restoring force but strong shear along the sum axis for *h* = 2 mV, leading to an overall strong accentuation of (co-)variability of E and I activities (Figure 2B). For larger inputs, both the feedforward and self-couplings grew with *h*, but the increasing quenching effect of self-couplings dominates the expanding effect of balanced amplification, leading generically to a pronounced net decrease in overall variability (Figure 1D, bottom; Figure 2C). For example, in the limit of slow noise, the summed E/I variance has a simple form that includes a term explicitly capturing the opposing effects of self couplings and balanced amplification: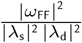. This term grows with the square of *ω*_FF_ but is divided by four powers of λ, indicating that self-couplings, if sufficiently strong, will dominate balanced amplification (for a derivation, and the more general case, see SI).

All the effects mentioned above were robust to changes in parameters, which we could show both through inspection of analytical formulae for activity variability and through numerical explorations of a thousand networks with randomly chosen parameters (SI).

### Variability quenching speeds up activity fluctuations

The growing restoring force also sped up the network dynamics, which was seen in the sharpening of the *V*_E/I_ autocorrelograms by large external inputs (Figure 1E). This was because the effective time constant with which fluctuations decay in the network is inversely proportional to the restoring force (Murphy and Miller, 2009). This speeding up was also reflected in the drop of LFP power at low frequencies (Figure 1F), in line with experimental data (Poulet and Petersen, 2008; Tan et al., 2014; Chen et al., 2014). At higher frequencies, this drop was over-compensated by the amplifying effect of the shear force and by the emergence of weak resonance, resulting in larger LFP power relative to the lowinput condition. Such a pattern of changes in the LFP has indeed been found in V1 of the awake macaque between evoked and spontaneous activity, although there was overall more power at high frequencies in both conditions than our model predicted (Tan et al., 2014). This may simply stem from an increased contribution of fast “spiking noise” at high firing rates in the cortex, which could not be captured by this population-level model but emerged naturally in a more detailed model of the same 2-population architecture using individual spiking neurons, as we show in the following.

### Variability reduction in a network of spiking neurons: impact of input noise correlations

In order to study variability in single neurons and at the level of spike counts, we implemented the two-population architecture of Figure 1A in a network of spiking neurons (Experimental Procedures). The network consisted of 4000 E neurons and 1000 I neurons, randomly connected with low probability, and with weights chosen such that the mean connectivity to an E or I neuron matched that to an E or I unit, respectively, in the reduced model. Each neuron emitted action potentials stochastically with an instantaneous rate given by Equation 2 (this additional stochasticity accounted for the effects of unmodelled fluctuations in synaptic inputs that occur on timescales faster than the 30 ms effective time resolution of our model). The external input to the network again included a constant term, *h*, and a noise term that was temporally correlated on a 50 ms timescale, and also spatially correlated with a uniform correlation across neurons, *ρ*. We systematically varied *h* and *ρ* to study their effects on the variability of responses in both spike count Fano factors and membrane potentials.

At the population level, for any level of input noise correlation ρ, the network behaved as predicted by the reduced model. Neurons fired irregularly (Figure 3A, C, top) with firing rates that grew superlinearly with small input *h* but sublinearly with stronger input (Figure 3E, top left). Moreover, fluctuations in E and I population activities were strongly synchronized (Figure 3A and C, bottom), and variability of these population-averaged rates and of the LFP (population-averaged *V*_m_) decreased with increasing *h* (although their absolute scale did depend on *ρ*; Figure 3A and C, bottom and B and D, top).

**Figure 3.**
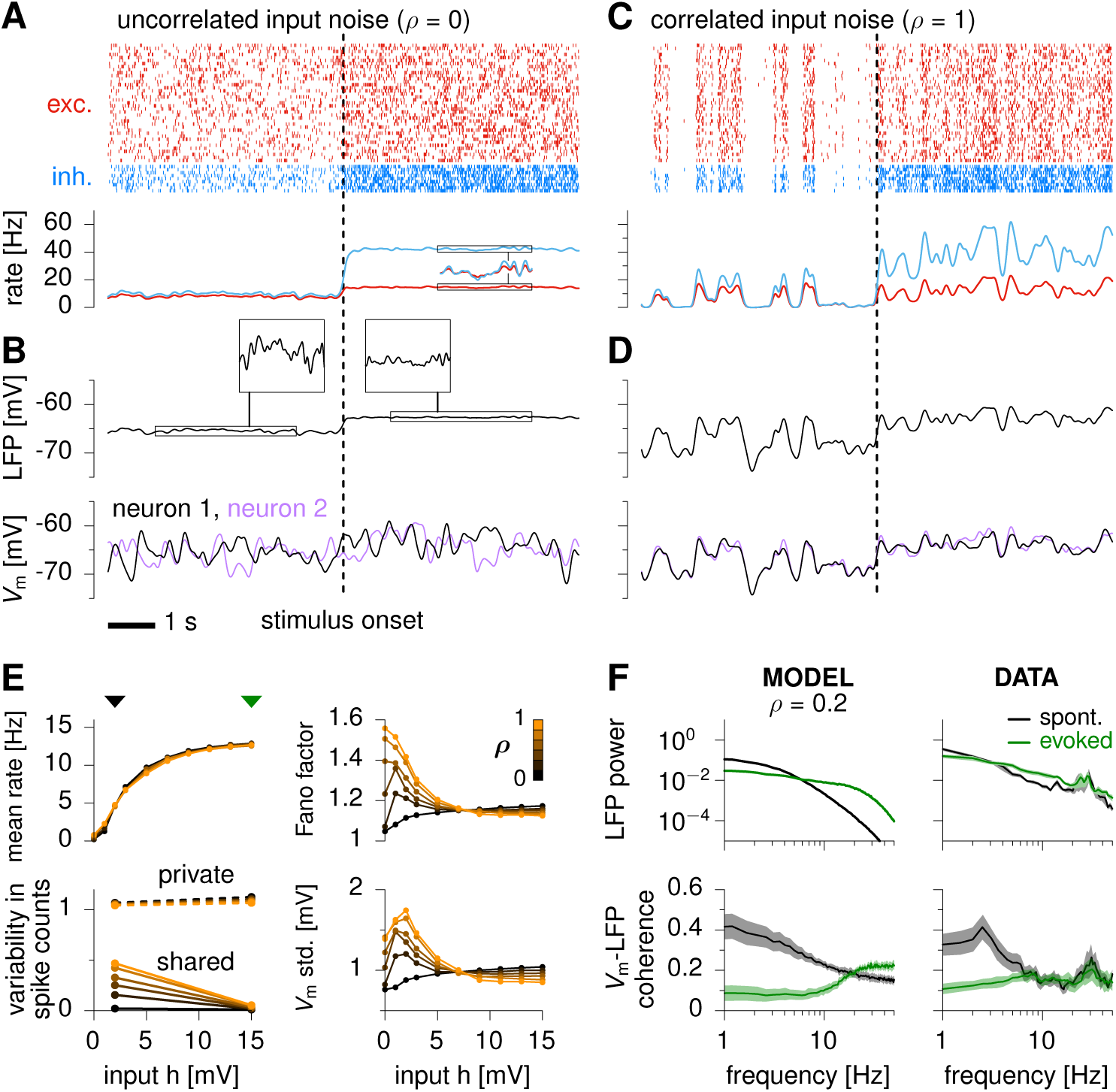
The modulation of variability in a randomly connected SSN. (**A**) Top: raster plot of spiking activity, for 40 (out of 4 000) excitatory neurons (red) and 10 (out of 1 000) inhibitory neurons (blue) when external input noise is private to each neuron (*ρ* = 0). The dashed vertical line marks the onset of stimulus, when *h* switches from 2 mV to 15 mV. Bottom: momentary population firing rate. The inset shows two overlaid segments on a magnified vertical scale. (**B**) Top: LFP (momentary population-averaged Vm). Insets magnify two segments with the same LFP scale, to visualise the relative drop in LFP variability following stimulus increase. Bottom: Vm of two randomly chosen units. (**C-D**) Same as (A-B) with external noisy inputs fully correlated across neurons (*ρ* = 1). (**E**) Mean firing rates (top left), private vs. shared parts of single-cell spike count variability as estimated by factor analysis (Bottom left), spike count Fano factors (top right) and Vm std. (bottom right) as a function of the external input *h*, for various values of the input correlation *ρ* (black to orange, 0 to 1 in steps of 0.2), and averaged over the E population. (**F**) Top: LFP power in spontaneous conditions and evoked conditions (black and green, respectively, cf. marks in panel C); Bottom: average (±s.e.m.) spectral coherence between single-cell *V*_m_ and the LFP; Left: model; Right: data from V1 of the awake monkey, reproduced from Tan et al., 2014. Firing rates, LFP, and Vm traces in panels A-E were smoothed with a Gaussian kernel of 50 ms width. In panel E, spikes were counted in 100 ms bins.

In contrast, variability reduction at the level of single neurons depended on the input noise correlation *ρ*. Single neuron variability quenching occured only when neurons shared part of their input noise, i.e. when *ρ* was sufficiently large (Figure 3B and D, bottom). For small *ρ*, individual *V*_m_ variances (Figure 3E, bottom right) had only a weak dependence on *h* (and, in fact, slightly grew with *h*, which could be explained by growing firing rates and hence increasing variance in synaptic input). With larger *ρ*, *V*_m_ variances decreased with increasing *h*, mirroring the quenching of LFP variability. In all cases, changes in *V*_m_ variability were directly reflected in Fano factors: strong *h* quenched spiking variability only for sufficiently large *ρ* (Figure 3E, top right). Indeed, Fano factors were well approximated by 1+*C*.var(*V*_m_) with some constant *C*, provided firing rates were not too small (Hennequin and Lengyel, *in prep*.). Note that changes in Fano factor with varying *ρ* could not be accounted for by changes in mean firing rates, which indeed had no dependence on *ρ* (overlapping colored lines in Figure 3E, top left).

The role of input correlations in variability quenching can be understood based on a decomposition of total spike count variability in each cell into a private noise term and a term that is shared with the other cells (Figure 3E, bottom left; here, shared and private variability are dimensionless and sum up to the average spike count Fano factor; see ‘Factor analysis’ section in Experimental Procedures). While the private noise term only depended on the private noise level in the input, the shared term depended on fluctuations in population activity. In turn, these population-wide fluctuations were fed by correlated input noise across neurons, and it was this shared variability that could be shaped by the interactions between E/I populations as predicted by the reduced model. Thus, when input correlations were small, single neuron variability was dominated by private noise with only minimal shared variability to be suppressed by increasing h (for *ρ* = 0, shared variability goes from 0.02 for *h* = 2 mV, to 0. 01 for *h* = 15 mV; cf. almost flat solid black line in Figure 3E, bottom left). As a consequence, no quenching of single-cell variability could occur (and in fact, since private variability grew with mean firing rate, single-neuron variability grew with *h*). LFP fluctuations were small, reflecting the small shared noise, because the uncorrelated private noise was effectively averaged out. In contrast, when input correlations were large, shared variability became substantial, leading to larger overall LFP fluctuations and larger reduction in singlecell variability by increasing input, *h*. This pattern of stimulus strength primarily modulating shared but not private variability is consistent with experimental findings in several cortical areas (Churchland et al., 2010).

Our model also accounted for the stimulus-induced modulation of the power spectrum and cross-coherence of LFP and single-cell *V*_m_ fluctuations, as observed in V1 of the awake monkey (Figure 3F; Tan et al., 2014). Consistent with the results obtained in the reduced rate model (Figure 1F), strong external input reduced the LFP power at low frequencies, and increased it at higher frequencies (Figure 3F, top left). This increase resulted from two effects. First, there was a small increase of LFP power at moderately high frequencies (Figure 1F), due to the input-induced increase in balanced amplification (shear force) outweighing the input-induced decrease in self-inhibition (restoring force) at those frequencies. Second the larger firing rates associated with strong inputs contributed additional fluctuations in synaptic drive on fast timescales due to stochastic spiking, thus increasing the relative variability in the LFP in higher frequency bands.

This asymmetric modulation of LFP power at low and high frequencies is also seen in the experimental data (Figure 3F, top right). Moreover, as strong input suppressed shared variability at low frequencies, the private noise in the activity of each neuron made a proportionately larger contribution to its overall variability at those frequencies, leading to a drop in *V*_m_-LFP coherence specifically at those frequencies where the suppression of population variability occurred, as seen in experiments (Figure 3F, bottom).

### stimulus-dependent suppression of variability in an SSN with structured connectivity

Neuronal recordings in area MT have shown that Fano factors drop at the onset of the stimulus (drifting gratings or plaids) in almost every neuron, which was well accounted for by the randomly connected networks we studied above. However, in the experiments, variability did not drop uniformly across cells, but exhibited non-trivial dependencies on stimulus tuning (Ponce-Alvarez et al., 2013; Lombardo et al., 2015). Similar effects were also observed in V1 of the anesthetized cat (Lin et al., 2015). This could not be explained by randomly connected architectures, and so we extended our model to include tuning-dependence in connectivity and input noise correlations.

We revisited the rate-based dynamics of Equation 1, now in an architecture in which the preferred stimulus of E/I neuron pairs varied systematically around a “ring” representing some angular stimulus variable, such as motion direction (Figure 4A; Experimental Procedures). The average input to a cell (either E or I) was composed of a constant baseline, which drove spontaneous activity in the network, and a term that depended on the angular distance between the stimulus direction and the preferred direction (PD) of the cell, and that scaled with stimulus strength, *c* (Figure 4C) – with c varying from 0 to 1 (increasing *c* represents increasing stimulus contrast). Input noise correlations depended on tuning differences (Experimental Procedures): cells with similar tuning received correlated inputs which in MT likely originate from upstream visual areas, such as V1, where activity fluctuations typically exhibit similar tuning-dependent correlations (Tsodyks et al., 1999; Kenet et al., 2003; Hansen et al., 2012; Ecker et al., 2010, 2014). Moreover, the strength of recurrent connections also depended on the difference in preferred direction between pre- and postsynaptic neurons, with the same tuning width for all connections whether excitatory or inhibitory (Figure 4B). This common tuning was based on the finding that, for the circular variable of orientation in cat V1, the excitation and inhibition that cells in layers 2-4 receive have the same tuning (Mariño et al., 2005; Martinez et al., 2002; Anderson et al., 2000). The model therefore differed from so-called “ring attractor” models which rely on similar topographic connectivity but with inhibition having wider tuning that excitation (Goldberg et al., 2004; Ben-Yishai et al., 1995; Ponce-Alvarez et al., 2013). This led to another important difference (discussed in Murphy and Miller, 2009): while attractor networks show sustained activity after stimulation even once the stimulus is removed, our network returned to baseline activity within a single membrane time constant (Figure 4D). As we show below, this dynamical regime is also characterized by fundamentally different patterns of response variability than attractor dynamics. Finally, to model spike count statistics, we assumed the same doubly-stochastic spiking mechanism as described above (Figure 3), but with spikes having no influence on the dynamics given by Equation 1 (we describe a fully spiking model later in Figure 8).

**Figure 4.**
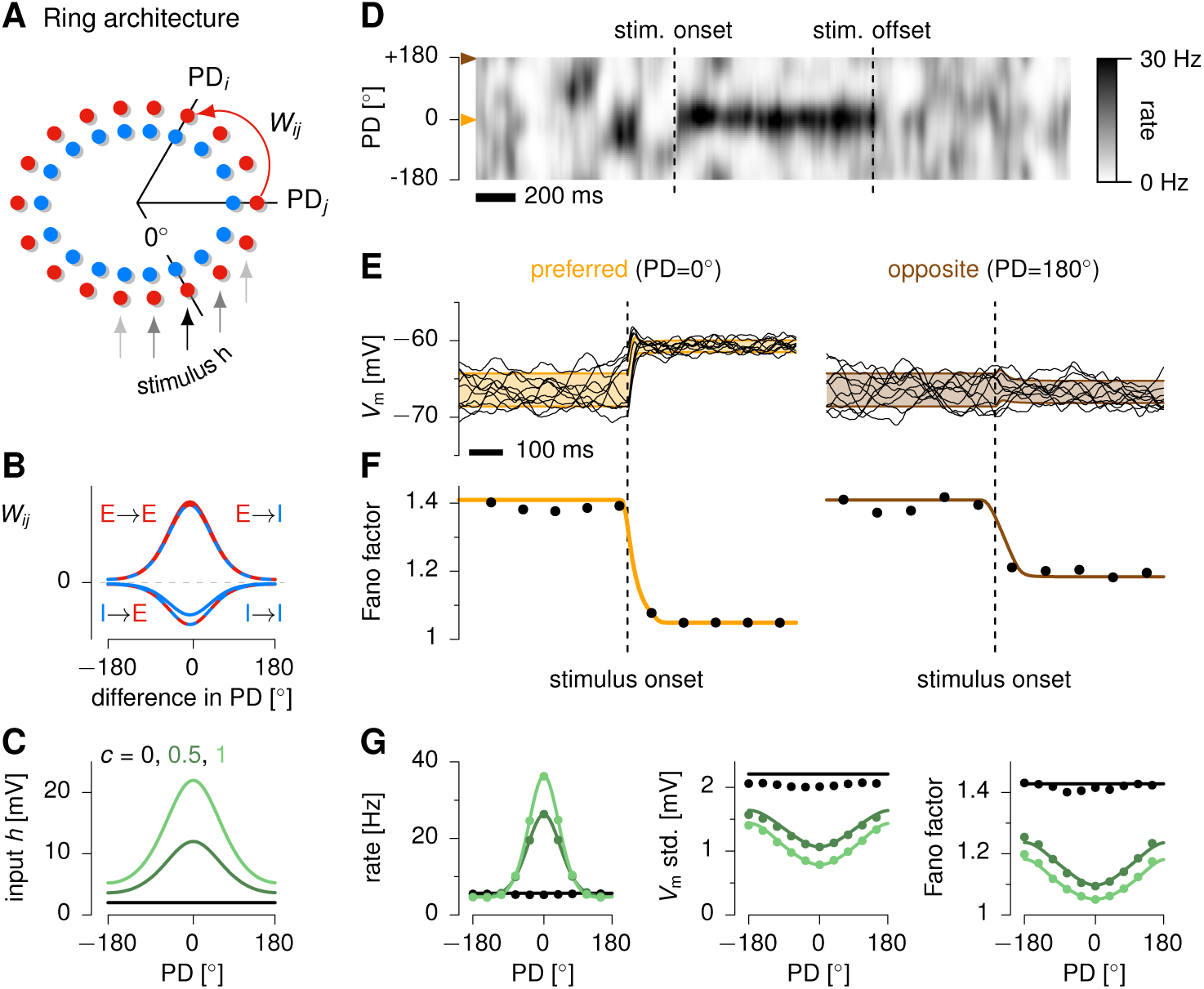
Across-trial variability in a ring SSN. (**A**) Schematics of the ring architecture. Excitatory and inhibitory neurons are laid out on a ring, their angular position ultimately determining their preferred stimulus (expressed here as preferred stimulus direction, PD) relative to the stimulus, assumed to be at 0° without loss of generality. (**B**) Synaptic connectivity follows circular Gaussian profiles with peak strengths that depend on the type of pre- and post-synaptic populations (excitatory or inhibitory). (**C**) Each neuron receives a constant input with a baseline (black line, *c* = 0), which drives spontaneous activity, and a tuned component with a bell-shaped dependence on the neuron’s preferred direction and proportional to stimulus strength *c* (dark and light green, *c* = 0.5 and *c* = 1 respectively). Neurons also receive spatially and temporally correlated noise, with spatial correlations that decrease with tuning difference (see Figure 5D).(**D**) Singletrial network activity (E cells), in response to a step increase and decrease in stimulus strength (going from *c* = 0 to *c* = 1 and back to *c* = 0). Neurons are arranged on the y-axis according to their preferred stimulus. (**E**) Reduction in membrane potential variability across 10 independent trials for an E cell tuned to the stimulus direction (left, corresponding to orange mark in D) or to the opposite direction (right, brown mark in D). (**F**) Reduction of spike count Fano factor following stimulus onset for the same two neurons as in (**E**). Spikes were counted in 100 ms time windows centered on the corresponding time points. (**G**) Mean firing rates (left), std. of voltage fluctuations (center) and Fano factors (right) as a function of the neuron’s preferred stimulus, at three different levels of stimulus strength (cf. panel C). Black lines in panel E and dots in panels F-G are based on numerical simulations over of 500 trials. Shaded areas in E and solid lines in F-G show analytical approximations (Hennequin and Lengyel, *in prep*.).

In the absence of visual input (*c* = 0), the input noise and mean baseline drove spatially patterned fluctuations in momentary firing rates around a few Hz (Figure 4D) with large across-trial variability in single-cell *V*_m_ (Figure 4E), implying super-Poisson variability in spike counts, i.e. Fano factors greater than 1 (Figure 4F). Visual stimulation drove a hill of network activity around the stimulus direction (Figure 4D), resulting in tuning curves of similar widths for different stimulation strengths (Figure 4G, left). Variability in both V_m_ and spike counts was strongly reduced compared to spontaneous conditions (Figure 4E-F), with variability reduction both for cells whose rate was increased by the stimulus (Figure 4E-F, left) and for those whose rate was unaffected (Figure 4E-F, right), as noted across many cortical areas (Churchland et al., 2010), but with a more pronounced reduction for cells whose preferred direction was close to the stimulus (Figure 4G). Notably, as in the randomly connected network of Figure 3, variability suppression in the ring model required finite spatial correlations in the input noise (Supplementary Figure S5).

### The effects of shear and restoring forces on bump dynamics explain structured patterns of variability

To understand the origin and mechanism of variability suppression in the ring architecture, we examined how recurrent interactions shaped the structure of *V*_m_ co-variability across the network. The most prominent feature of population activity was a “bump” of high *V*_m_ in the cells with preferred directions near the stimulus direction, and lower activity in the surround (Figure 5A). Accordingly, most of the shared variability (∼ 90%; Figure S4) arose from the variability in the location *μ*, amplitude *a* and width *σ* of this bump (Figure 5A and C). Each of these small transformations resulted in a pattern of momentary deviation of network activity from the propotypical bump (Figure 5B, right). In turn, the momentary fluctuations caused by these ongoing transformations (Figure 5C) contributed distinct spatial templates of covariance (Figure 5D). For example, sideways motion of the bump increased the firing rates of all the cells with preferred directions on one side of the stimulus direction, and decreased firing rates for all cells on the other side (Figure 5B, top). This resulted in positive correlations between cells with preferred directions on the same side of the stimulus direction, and negative correlations for cells on opposite sides (Figure 5D, *μ*-template; Moreno-Bote et al., 2014). Fluctuations in bump amplitude generated modest positive covariances that were somewhat greater between cells tuned near the stimulus direction (Figure 5D, middle left). In contrast, fluctuations in the width of the bump generated large positive covariances, especially between cells tuned near the opposite direction (Figure 5D, bottom left). As the nonlinear interactions among neurons result in strong normalization of overall activity in the dynamical regime of our network (Ahmadian et al., 2013; Rubin et al., 2015), fluctuations in amplitude and width were strongly (negatively) correlated, which contributed a distinct pattern of covariance: strong negative correlations for all pairs but those tuned to the stimulus (blue template in Figure 5D, left).

**Figure 5.**
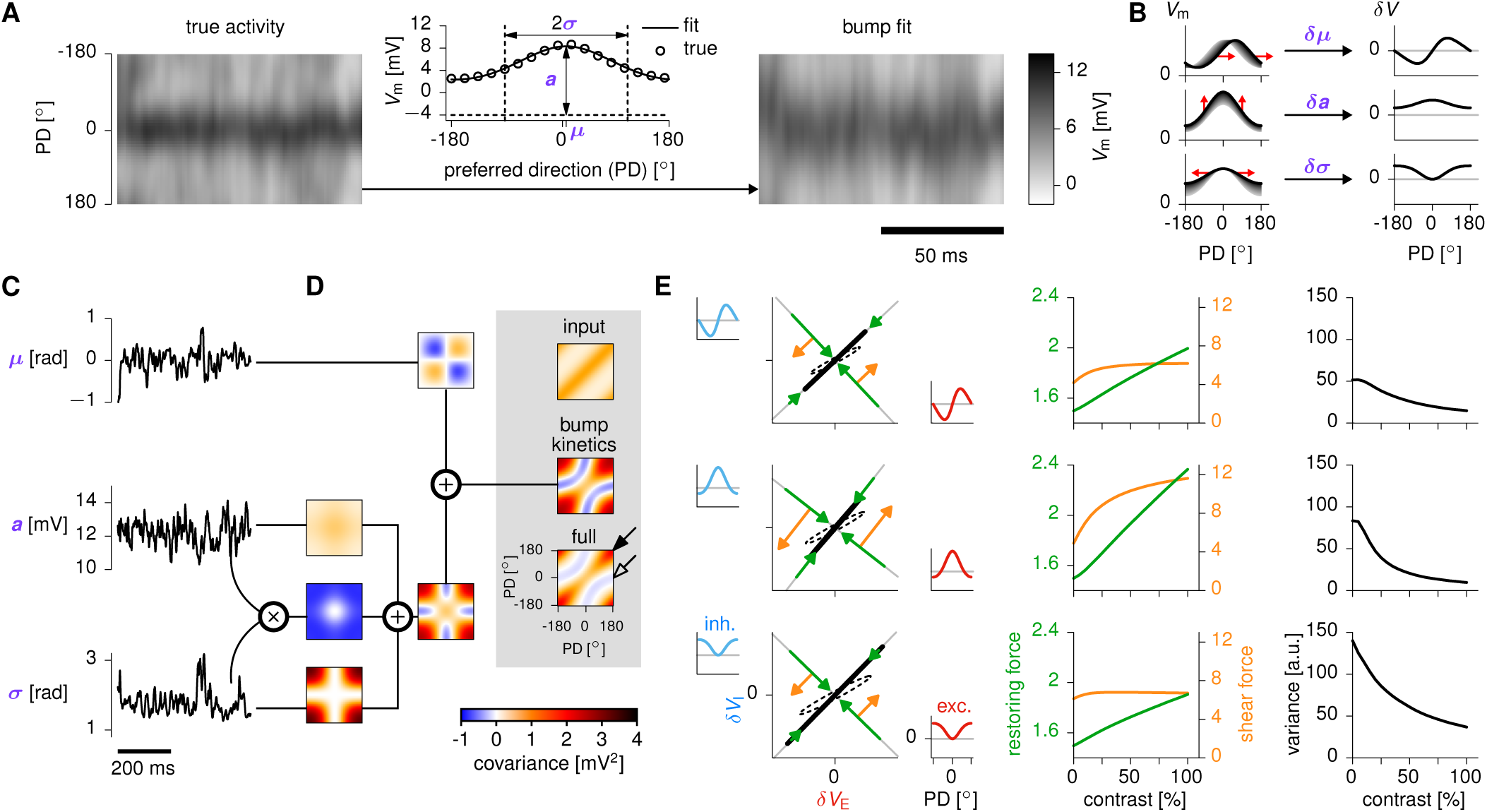
Differential reduction of variability in the three principal dimensions of bump kinetics. (**A**) 100 ms-long sample of *V*_m_ fluctuations across the network in evoked conditions (left, “true activity”, *c* = 1), to which we fitted a circular-Gaussian function *V*_i_(*t*) = – *a*_0_ + *a*(*t*)exp[(cos(*θ*; – *μ*(*t*)) – 1)/*σ*(*t*^2^] across the excitatory population in each time step (center). This fit captured most of the variability in *V*_m_ (right). (**B**) The three principal modes of bump kinetics: small changes (red arrows) in location (top), amplitude (middle) and width (bottom) of the activity bump result in the hill of network activity deviating from the prototypical bump (gray shadings). Plots on the right show how the activity of each neuron changes due to these modes of bump kinetics. (**C**) Time series of *μ*, *a* and *σ* extracted from the fit. (**D**) Ongoing fluctuations in each of the three bump parameters contribute a template matrix of *V*_m_ covariances among E cells (color maps), obtained from (the outer product of) the differential patterns on the right of panel B. The strong (anti-)correlation between *a* and *σ* contribute a fourth effective template. These templates sum up to a total covariance matrix (“bump kinetics”), which captures the key qualitative features of the full *V*_m_ covariance matrix (“full”). The covariance matrix of the input noise (“input”) is also shown above for reference. (See text for arrows.) (**E**) Left: three planes of spatially patterned E/I activity in which the recurrent dynamics of the network approximately decoupled (SI), corresponding approximately to the three modes of bump kinetics (compare axis insets to the differential patterns in panel B, right). Arrows show forces (orange: shear, green: restoring), ellipses show output covariances due to single-cell leak only (dashed) or full recurrent dynamics (solid), as in Figure 2B-C. Middle: dependence of forces on stimulus strength. Green curves show the average of the self-inhibitory couplings, |λ_d_| and |λ_5_|(λ_d_ and λ_s_ are shown individually in Supplementary Figure S4). Orange curves show the feedforward (shear force) coupling, |*ω*_FF_|. Right: the variance in the E population (projection of the solid ellipse onto the x-axis in each plane) as a function of the input strength *c*.

Taken together, the ongoing jitter in bump location, amplitude and width contributed a highly structured pattern of response covariances, which accounted for most of the structure in the full covariance matrix of the network (Figure 5D, compare “bump kinetics” with “full”). In particular, bump kinetics explained the comparatively stronger reduction of *V*_m_ variance for cells tuned near 0° (compare Figure 4G, middle, with the diagonal of the full covariance matrix indicated by the filled arrow in Figure 5D). Moreover, the recurrent dynamics generated negative correlations in the *V*_m_ fluctuations of cells with opposite tuning, despite such pairs receiving positively correlated inputs (Figure 5D, “input” vs. “bump kinetics”, secondary diagonal with open arrow).

Bump kinetics were not only useful to phenomenologically capture most of the covariability in the network, but they were also identified approximately as the most accurate lowdimensional summary of the recurrent dynamics by formal reduction techniques (SI). Reducing the dynamics of our model to these three motion modes revealed that the same forces that shaped variability in the two-population architecture also explained the more detailed patterns of variability reduction in the ring architecture (Figure 5E). However, while the two-population model only involved forces in a single plane describing population-averaged E and I activities (Figure 2B–C), the ring architecture induced forces in three different such planes involving three pairs of activity patterns in the E and I populations (Figure 5E, insets along plane axes) that corresponded almost exactly to the three modes of bump kinetics (Figure 5B; the “bump amplitude pattern” differs slightly, due to the requirement that it be orthogonal to the other two patterns).

As fluctuations in the external input were correlated among similarly tuned neurons irrespective of their E/I nature (“input” covariance matrix in Figure 5D), they instated correlated baseline *V*_m_ fluctuations in the E and I populations in each of the three planes where most variability was confined (Figure 5E, elongated dashed ellipses, obtained by neglecting the effect of recurrent connectivity). As in the two population model, the recurrent interactions modified both the restoring and shear forces (green and orange arrows in Figure 5E), which in turn amplified baseline *V*_m_ variability (solid ellipses). Patterns of momentary E/I imbalance (e.g. resulting from the E bump having moved more than the I bump) were strongly amplified into balanced patterns (Figure 5E, orange arrows, or “shear force”), and restoring forces acted to quench both imbalanced and balanced fluctuations (green arrows). These forces depended on the effective connectivity, which in turn depended on stimulus strength *c* (Figure 5E, center) such that restoring forces increased steadily with *c*, while shear forces saturated already at low values of *c* – just as seen in the two-population model (Figure 2E). Overall, restoring forces became increasingly dominant over shear forces, resulting in a reduction of variability in each of the three modes of bump kinetics with increasing *c* (Figure 5E, right). This reduction occured at different rates in the three modes, such that at high *c* variability was mostly caused by fluctuations in bump width, thus explaining the U-shape of *V*_m_ variance (Figure 4G, middle). Note that this yields interesting predictions for changes in the tuning of variances and covariances across the full range of stimulus strengths: in essence, a smooth morphing from the spontaneous covariance, for very low contrast, to the high-contrast covariance (Figure 7A, right). Moreover, as variability quenching occured predominantly in these three spatially very smooth activity modes, suppression of variability in single neurons could only occur provided these modes explained a sufficiently large fraction of the total network variance. This in turn required the input noise to contain spatially smooth correlations (Supplementary Figure S5).

### Differences between the SSN and attractor models

For a direct comparison of the SSN with attractor dynamics, we implemented a canonical model of attractor dynamics that have also been suggested to account for stimulus-modulated changes in variability (Ponce-Alvarez et al., 2013), and matched it to our model such that it produced similar tuning curves and overall levels of variability (Supplementary Figure S6). In contrast with the richer patterns of variability generated by our model, attractor dynamics showed a more limited repertoire, dominated solely by sideways motion of the bump. Moreover, restoring forces induced by attractor dynamics dominated over the shear forces at all stimulus strengths. As a result, the sign of membrane potential covariances depended on whether two cells had their preferred directions on the same side of the stimulus direction (Figure 5D, *μ*-template), but not otherwise on the difference between their preferred directions.

These differences in membrane potential covariances also carried over to spike count noise correlations that are experimentally more readily accessible. Most prominently, the attractor network predicted large negative correlations for cells tuned to opposite directions, whereas the SSN predicted predominantly positive correlations with only very weak negative correlations (Figure 6A-B). We note that it might seem trivial to eliminate negative correlations in the attractor network by invoking an additional (potentially extrinsic) mechanism that adds a single source of shared variability across neurons. This would result in a uniform (possibly stimulus strength-dependent) positive offset to all correlations (Lin et al., 2015). However, the two models also exhibited differences that would not be explained even by this additional mechanism. Specifically, in the SSN, spike count correlations for pairs with a fixed difference in preferred directions (fixed ΔPD) depended only weakly on the stimulus direction (in Figure 6A-B, lines parallel to the lower-left to upper-right diagonal represent pairs with a fixed ΔPD, while a change in stimulus direction for a given pair corresponds to movement along such a line; also note similarities between the three panels in Figure 6E). In contrast, in the attractor network at high stimulus strength, spike-count correlations for a pair of fixed ΔPD can depend strongly on stimulus direction (Figure 6B, F). Moreover, while both models predicted noise correlations to generally decrease with ΔPD, stimulus strength simply scaled this decrease in the SSN approximately uniformly (Figure 6E), but interacted with ΔPD in more complex ways in the attractor network, such that correlations could change proportionately more or less for different cell pairs (Figure 6F).

**Figure 6.**
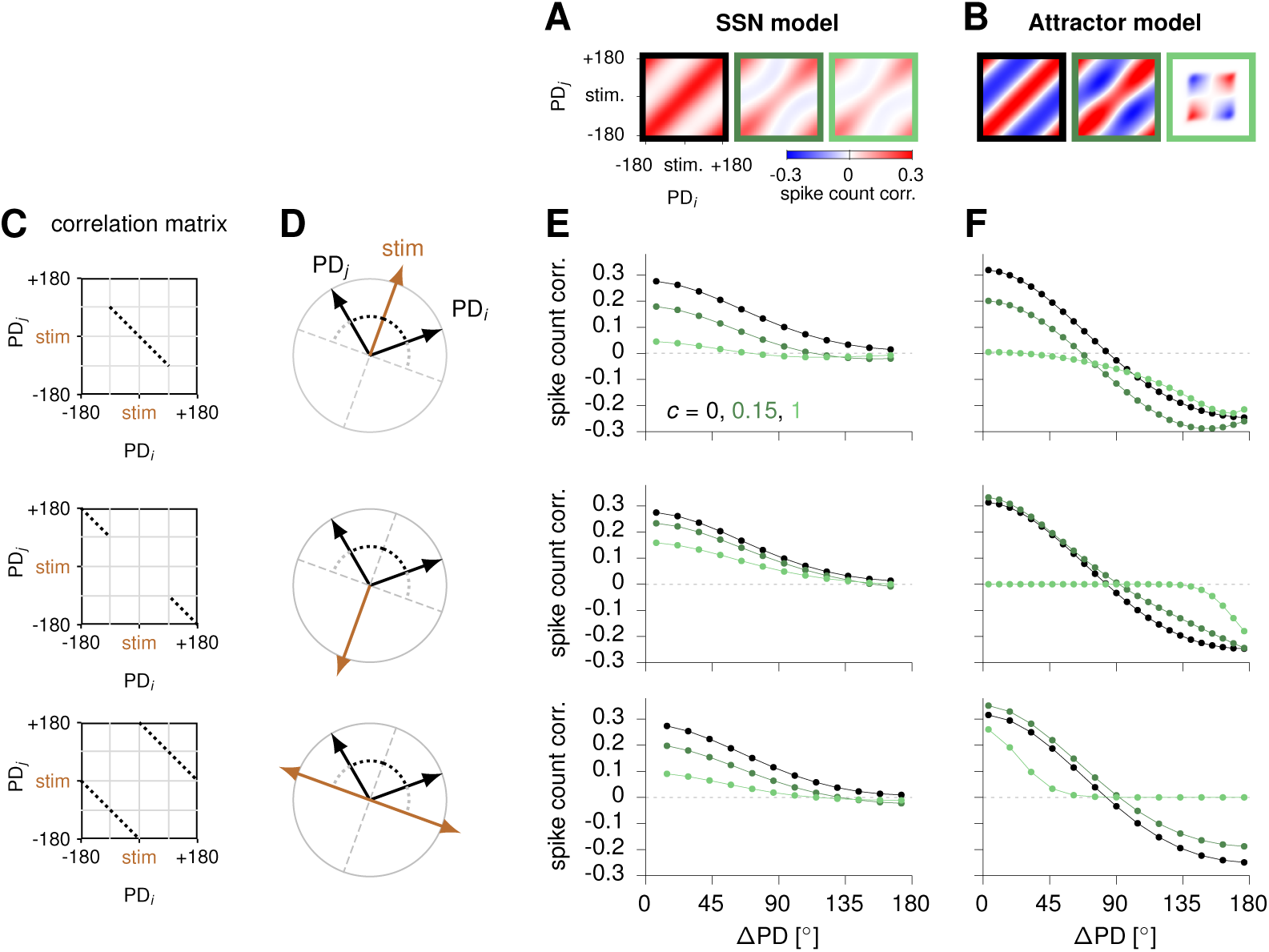
The SSN and ring attractor network make distinct predictions for spike count noise correlations. (**A**) Spike count correlation matrices in the SSN, for three values of stimulus strength (border color black: *c* = 0; dark green: *c* = 0.15; light green: *c* = 1). X- and y-axes of each matrix are preferred directions (PDs) of two cells, relative to stimulus direction taken equal to 0. (**B**) Same as (A), for the ring attractor network.(**C**) Spike count correlations in the SSN and attractor network most strongly differ along particular “cross-sections” of the correlation matrices (dotted line segments).(**D**) The segments shown in (C) correspond to scenarios in which the stimulus direction exactly bisects the (smaller) angle between the preferred directions of the two recorded cells (top), or is opposite (middle) or orthogonal (bottom) to this direction. Each difference between preferred directions, corresponds to a specific position on the dotted segments in (C).(**E**) Spike count correlations as a function of ΔPD, along the segments shown in the corresponding matrices in (C) at different stimulus strengths (colors as in A-B). (**F**) Same as (E), for the ring attractor network.

### Comparison to variability of responses in MT

In our ring SSN model of directional tuning, the most robust effect concerning variability modulation by stimuli is a comparatively stronger drop in Fano factor and *V*_m_ variance in the neurons most strongly driven by the stimulus. Although this “U shape” of variability quenching was also recorded in area MT of the awake macaque for some types of stimuli (namely coherent plaids; cf. top panel of Figure 1B in Ponce-Alvarez et al., 2013), other sets of stimuli instead resulted in an M-shaped profile of Fano factor reduction (see also Lombardo et al., 2015). Specifically, stimulus onset quenched variability more strongly in cells tuned to either the stimulus direction or the opposite one, compared to neurons tuned to the orthogonal directions (Figure 7C, center). A similar M shape was apparent for spike count correlations between similarly tuned neurons, as a function of their (common) preferred direction (Figure 7C, right).

**Figure 7.**
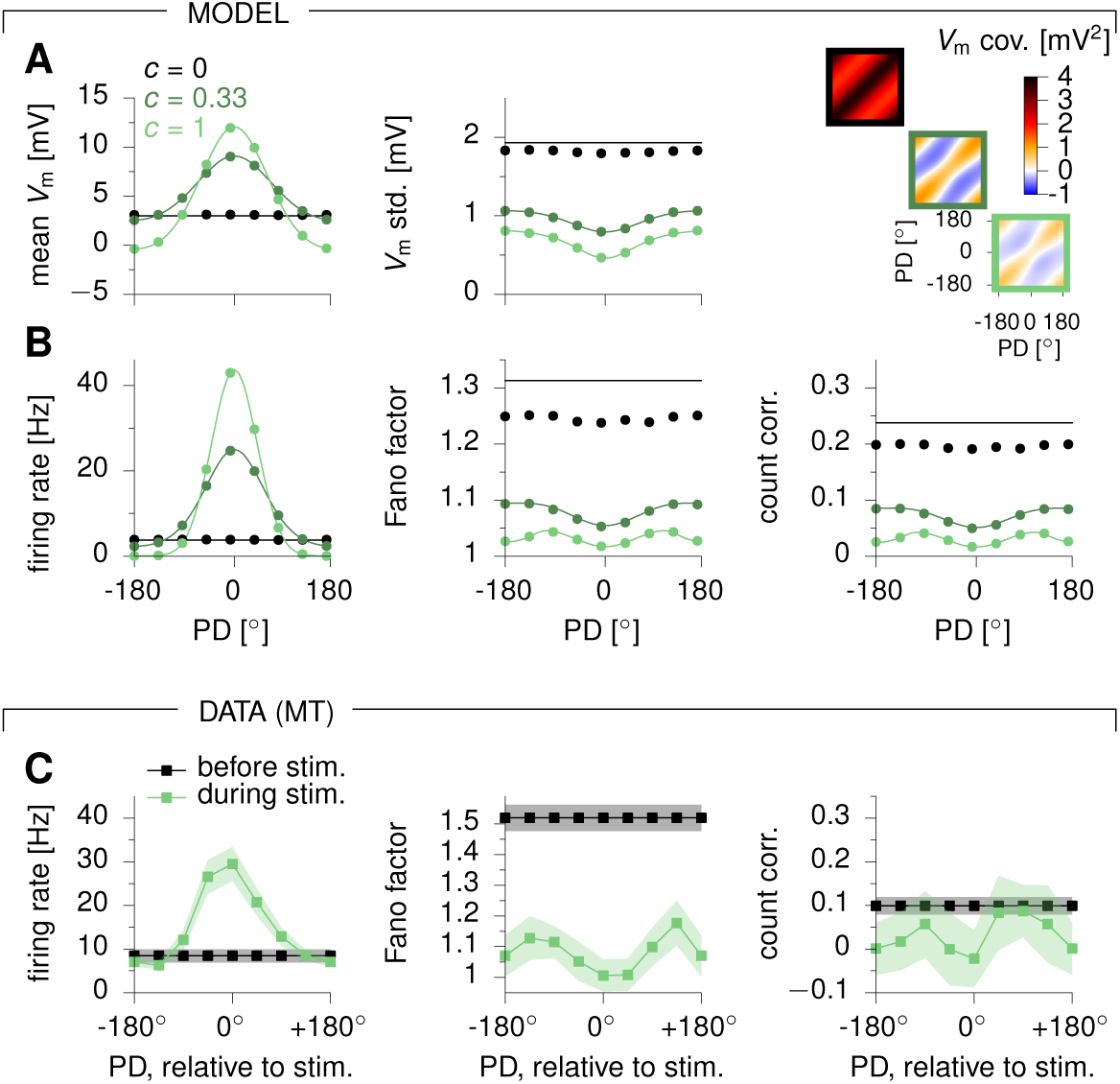
A ring SSN accounts for the stimulus dependence of across-trial variability in area MT. (**A**) Vm mean (left) and std. (center) as a function of the model neuron’s preferred direction, for increasing values of stimulus strength *c*. The full Vm covariance matrices are shown on the right for the E population, box color indicating *c*. (**B**) Mean firing rates (left), spike count Fano factors (center) and spike count correlations between similarly tuned neurons (right), as a function of the neuron’s preferred direction. (**C**) Experimental data (awake monkey MT) adapted from Ponce-Alvarez et al. (2013), with average firing rates (left), average Fano factors (center) and average spike count correlations among similarly tuned cells (right), as a function of the the cell’s preferred direction (PD, relative to stimulus at 0° shown for spontaneous (pre-stimulus, black) and evoked (*c* = 1 stimulus, green) activity periods. Error bars denote s.e.m. Dots in panels A-B were obtained from 400 s epochs of simulated stationary activity, and denote averages among cells with similar tuning preferences (PD diference<18°);solid lines show analytical approximations (Hennequin and Lengyel, *in prep*.). In panels B-C, spikes were counted in 100 ms bins.

We found that our model could also exhibit such an M-shaped modulation of both Fano factors and pairwise correlations at high stimulus strength (Figure 7B). This occurred when the network was set up such that cells tuned to the opposite direction became near-silent (Figure 7B, left), which typically required the tuning of the external input to be spatially as narrow as, or narrower than, that of the recurrent connectivity. In this case, the mean *V*_m_ of cells tuned to the opposite direction became comparable to, or smaller than, the rectification threshold *V*_rest_ in Equation 2, such that nearly half of their membrane potential fluctuations did not pass the rectification and thus had no effect on momentary firing rate fluctuations. Even the part of membrane potential fluctuations which passed the rectification threshold were diminished in the output by the small gain of the power-law neuronal nonlinearity close to its threshold. Thus, although the membrane potential fluctuations were larger for these cells than for orthogonally tuned neurons (Figure 7A, center), a substantial fraction of these fluctuations dissipated below threshold or were diminished by the small neuronal gain, yielding a lower firing rate variance. In fact, this loss of firing rate variance more than overcame the effect of dividing by very small firing rates in computing Fano factors for these neurons (Figure 7B, center). A similar nonlinear effect caused spike count correlations among similarly tuned neurons to exhibit an M-shape modulation at high stimulus strength (Figure 7B, right).

All our main results were reproduced in a sparsely connected spiking model of area MT, similar to that of Figure 3 but with an underlying ring architecture as in Figure 4 (Experimental Procedures). Single neurons fired action potentials asynchronously and irregularly during both spontaneous and evoked conditions (Figure 8A). Mean firing rates had an approximately invariant tuning to stimulus direction across stimulus strengths *c* (Figure 8D), and saturated strongly at large values of *c* (not shown). Moreover, both membrane potential variances and Fano factors decreased at stimulus onset (Figure 8B-C), and this drop in variability was also tuned, thus reproducing the M-shaped modulation of Fano factors and spike count correlations of the rate model (Figure 8E-G). Consistent with the analyses of bump kinetics and of the randomly connected spiking network, factor analysis revealed that stimulus quenched shared, but not private, variability in single neurons (Figure 8H). This indicated that the insights we obtained from studying simplified network architectures about the conditions for observing variability quenching in single neurons also applied to the ring architecture.

**Figure 8.**
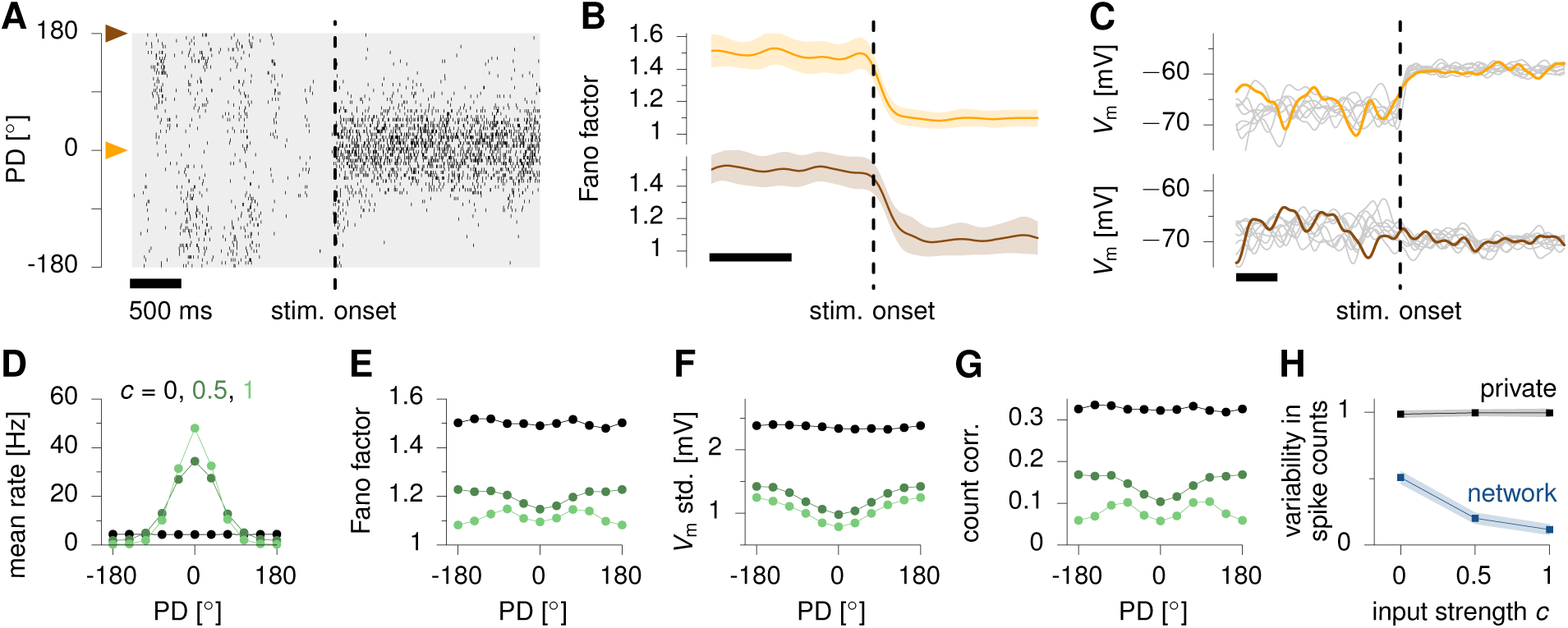
Variability suppression in a spiking network model of area MT. (**A**) Raster plot of spiking activity in excitatory neurons, arranged vertically according to preferred motion direction (PD). Activity is shown for 4 s around stimulus onset (dashed vertical line). (**B**) Fano factor time course for two E cells respectively tuned to the stimulus direction (orange mark in panel A) and to the opposite direction (brown mark), obtained from 1000 independent trials. (**C**) Single-trial Vm traces for the two cells shown in (B). One trial stands out in color, and 9 other trials are shown in gray to illustrate reduction of variability both within- and across trials. (**D–G**) Mean firing rates (D), Fano factors (E), Vm std. (F) and spike count correlations between similarly tuned cells (G), as a function of preferred direction (PD), at 3 different levels of stimulus strength c (color-coded as indicated in D). (**H**) Factor analysis performed on spike counts (Experimental Procedures), separating the private (black) from the shared (blue, “network”) contributions to spike count variability in every neuron. Shown here are mean private and shared variability across neurons, ± std across a subset of 200 randomly chosen neurons in the excitatory population. In panels E, G and H, spikes were counted in 100 ms bins. All statistics were estimated from 400 seconds of stationary simulated activity for each value of *c*, and averaged among cells with similar tuning preferences (PD difference<18°)In panels C and F, *V*_m_ fluctuations were first smoothed with a 50 ms Gaussian kernel.

## Discussion

We studied the modulation of variability in a stochastic, nonlinear model of cortical circuit dynamics. We focussed on a simple circuit motif that captured the essence of cortical networks: noisy excitatory and inhibitory populations interacting in a recurrent but stable way despite expansive single-neuron nonlinearities. This stochastic stabilized supralinear network (SSN) reproduced key aspects of variability in the cortex. During spontaneous activity, i.e. for weak external inputs, model neurons showed large and relatively slow synchronous fluctuations in their membrane potentials, which were quenched and decorrelated by stronger stimuli. The model thus explains and unifies a large body of experimental observations made in diverse systems under various conditions (Churchland et al., 2006, 2010; Finn et al., 2007; Poulet and Petersen, 2008; Gentet et al., 2010; Poulet et al., 2012; Tan et al., 2014; Chen et al., 2014). Moreover, the drop in variability was tuned to specific stimulus features in a model of area MT, also capturing recent experimental findings (Ponce-Alvarez et al., 2013; Lin et al., 2015; Lombardo et al., 2015). The SSN also captures ubiquitous phenomena involving nonlinear response summation to multiple stimuli, including normalization, surround suppression, and their dependencies on stimulus contrast (Rubin et al., 2015). Together these results suggest that the “loosely balanced” SSN captures key elements of the operating regime of sensory cortex.

Our analysis relied on the reduction of the complex mesh of recurrent, feedback-driven interactions among multiple neuronal populations into two types of effective connections among activity patterns (Murphy and Miller, 2009): a self-connection that, when including the single-cell leak (the tendency of isolated neurons to return to rest) as well as network synapses, must be inhibitory in a network that has stable steady-state responses to steady input, and which thus constitutes a “restoring force” that contains or quenches variability; and a feedforward pattern of connections between activity patterns that instantiates “balanced amplification”, amplifying small momentary disturbances of the E/I balance into large but balanced responses, and which can be thought of as a “shear force” boosting response variability. Crucially, this effective network connectivity depends on the mean firing rates of the E and I cells through the nonlinear response properties of the single neurons, and therefore depends on the strength of the external input. Balanced amplification typically dominates during spontaneous activity (i.e. for small to moderate inputs), increasing variability relative to that of isolated cells with the same external input; while for larger inputs, inhibitory self-connections become dominant, quenching this spontaneous variability (whether the peak of variability lies at spontaneous or at external input levels somewhat below or above those of spontaneous activity remains unclear). Importantly, these insights carried over to the higher dimensional, structured ring architecture used to model MT responses, providing the logical link between the network’s bumps of population activity in response to tuned inputs and the resulting structured, contrast-dependent patterns of variability it generated.

The SSN reproduces (Ahmadian et al., 2013; Rubin et al., 2015) much of the phenomenology of the “normalization model” of cortical responses (Carandini and Heeger, 2012) and provides a circuit substrate for it. In the normalization model and the SSN, responses to multiple stimuli add sublinearly, and as one stimulus becomes stronger than another, the response to their simultaneous presentation becomes “winner-take-all”, more and more dominated by the response to the stronger stimulus alone. This behavior predicts some aspects of variability suppression: a stronger mean input drive relative to the noise input leads to greater suppression of the noise’s contribution to the neuron’s response.

### Further factors modulating variability

We analyzed variability modulation solely as arising from intrinsic network interactions, but other factors may also contribute (Doiron et al., 2016). External inputs may be modulated; for example, the drop with contrast in LGN Fano factors has been argued to underlie *V*_m_ variability decreases in V1 simple cells (Sadagopan and Ferster, 2012; but see Malina et al., 2016). However, since high contrast stimuli also cause firing rates to increase in LGN, the total variance of LGN-to-V1 inputs (scaling with the product of the LGN Fano factor and mean rate) is modulated far less by contrast. This provides some justification for our model choice that input variance did not scale with contrast. Cellular factors may also modulate variability. For example, inhibitory reversal potential or spike threshold may set boundaries limiting voltage fluctuations, which would more strongly limit voltage fluctuations in more hyperpolarized or more depolarized states respectively; conductance increases will reduce voltage fluctuations; and dendritic spikes may contribute more to voltage fluctuations in some states than others (Stuart and Spruston, 2015). A joint treatment of external input, cellular, and recurrent effects may be needed to explain, for example, why *V*_m_ variability appears strongest near the preferred stimulus in anaesthetized cat V1 (Finn et al., 2007), or why overall V_m_ variability grows with visual stimulation in some neurons of awake macaque V1 (Tan et al., 2014).

Neuromodulators (and presumably anesthetics) can alter the input/output gain of single neurons as well as synaptic efficacies (Disney et al., 2007; Marder, 2012), yielding changes in effective connectivity that may in turn explain brain statedependent changes in cortical variability (Poulet and Petersen, 2008; Ecker et al., 2014; Lin et al., 2015; Mochol et al., 2015; Lombardo et al., 2015). Our approach, deriving changes in variability directly from changes in effective connectivity, offers a framework for also understanding these forms of variability modulation. Modifications of actual synaptic connections also alter effective connectivity, so our efforts are complementary to those of previous studies that focussed on the consequences for correlations of different anatomical connectivity patterns (Kriener et al., 2008; Tetzlaff et al., 2012; Ostojic, 2014; Hennequin et al., 2014b).

### The dynamical regime of cortical activity

We found that variability quenching in the stochastic SSN robustly occurred as the input pushed the dynamics to stronger and stronger inhibitory dominance. Consistent with this, with increasing strength of external input the ratio of inhibition to recurrent excitation received by SSN cells increases (Rubin et al., 2015), as also observed in layers 2/3 of mouse S1, in recordings in non-optogenetically-excited pyramidal cells, with increasingly strong optogenetic excitation of other pyramidal cells (Shao et al., 2013). This distinguishes the SSN from the balanced network (van Vreeswijk and Sompolinsky, 1998), for which this ratio would be fixed for a given pattern of external input to cells, regardless of the strength of activation. The two models are also distinguished by the nonlinear behaviors seen in SSN and cortex but not in the balanced network (discussed in Introduction). Finally, the balanced network predicts that external input alone is very much larger than the net input (recurrent plus external). In contrast, the SSN allows external and net input to be comparable, as observed in intracellular recordings in V1 layer 4 when the external thalamic input is revealed by suppressing cortical spiking (Ferster et al., 1996; Chung and Ferster, 1998; Lien and Scanziani, 2013; Li et al., 2013).

Two proposals have been made previously to explain quenching of variability by a stimulus: a stimulus may quench multi-attractor dynamics to create single-attractor dynamics (Blumenfeld et al., 2006; Litwin-Kumar and Doiron, 2012; Deco and Hugues, 2012; Ponce-Alvarez et al., 2013; Doiron and Litwin-Kumar, 2014; Mochol et al., 2015); and a stimulus may quench chaotic dynamics to produce non-chaotic dynamics (Molgedey et al., 1992; Bertschinger and Natschlger, 2004; Sussillo and Abbott, 2009; Rajan et al., 2010; Laje and Buonomano, 2013). In a ring architecture, our model differs from multi-attractor dynamics in two fundamental ways. First, attractor dynamics yields patterns of network variability originating almost exclusively from sideways motion of the activity bump (Supplementary Figure S6), leading to an M-shaped profile of Fano factor suppression. Although our model could also reproduce this M shape (Figures 7 and 8), it also exhibited substantial fluctuations in bump amplitude and width, producing a richer – yet still low-dimensional – basis of variability patterns which more typically combined to give Fano factors profiles a “U” shape (Figure 4). Indeed, coherent plaids or random dot stimuli in the macaque (Ponce-Alvarez et al., 2013; Lombardo et al., 2015) as well as in the marmoset (Sam Solomon, personal communication) result in a pronounced U-shaped modulation of Fano factors in MT. Our analysis suggested that the SSN can produce either M- or U-shaped modulations depending on the tuning width of inputs relative to that of connectivity, but that in both cases membrane potential variability will still have a U-shaped profile (Figures 7 and 8), which could be tested in future experiments. Second, patterns of bump motion also led to very different patterns of covariances and correlations across the population in the two models (Figure 5D). For strong input, attractor dynamics exclusively predict negative correlations for all cell pairs whose preferred stimuli are on opposite sides of the stimulus (Figure 6B,F; top left and bottom right quadrants of the *μ* covariance matrix in Figure 5D and Supplementary Figure S6; Ponce-Alvarez et al., 2013; Wimmer et al., 2014), while the SSN predicts that cells with similar tuning will be positively correlated even if the stimulus lies between their preferred stimuli (Figure 6A,E; Figure 5D, center and corners of the full covariance matrix). So far, correlations have not been reported as parametric functions of both the stimulus and the tuning differences of cells, or only in the context of attentional manipulations (Cohen and Newsome, 2008), leaving these predictions to be tested in future experiments. Thus, our work suggests a principled approach to use data on cortical variability to identify the dynamical regime in which the cortex operates.

More generally, our results also propose a very different dynamical regime underlying variability quenching than the multi-attractor or chaos-suppression models. The SSN differs from these in exhibiting a single stable state in all conditions – spontaneous, weakly-driven, strongly-driven – whereas the others show this only when strongly driven. Furthermore, quenching of variability and correlations in the SSN is highly robust, arising from two basic properties of cortical circuits: inhibitory stabilization of strong excitatory feedback (Tsodyks et al., 1997; Ozeki et al., 2009), and supralinear input/output functions in single neurons (Priebe and Ferster, 2008). In contrast, models of multi-attractor or chaotic dynamics can either account only for the modulation of average pairwise correlations (Mochol et al., 2015), or else require considerable fine tuning of connections (Litwin-Kumar and Doiron, 2012; Ponce-Alvarez et al., 2013) to account for more detailed correlation patterns. Moreover, as studied thus far they typically ignore Dale’s law (the separation of E and I neurons) and its consequences for variability, e.g. balanced amplification (Rajan et al., 2010; Ponce-Alvarez et al., 2013; Mochol et al., 2015) (but see Harish and Hansel, 2015; Kadmon and Sompolinsky, 2015).

Other differences of dynamical regime suggest further experimental tests. Mechanisms of chaos control typically lead to quenching of across-trial variability at stimulus onset, but not within-trial variability across time (Sussillo and Abbott, 2009; Rajan et al., 2010; Laje and Buonomano, 2013), as could be assayed by measures of variability in sliding windows across time. Both the SSN and multi-attractor models predict quenching of both forms of variability. In chaotic models, the transition from high- to low-variability is sudden with increasing external input strength (Rajan et al., 2010), while the transition in the SSN will be, and in multiattractor models may be, gradual. In the high-variability spontaneous state and for weakly-driven states (i.e. for a low-contrast stimulus), the chaotic and multiattractor scenarios both predict slow dynamics (relative to cellular or synaptic time constants), measurable as long auto-correlation times for neural activity (Sompolinsky et al., 1988; Sussillo and Abbott, 2009; Rajan et al., 2010; Laje and Buonomano, 2013) and as slow responses to stimulus changes. Dynamics in these scenarios may become fast in the high-input, low-variability state. In contrast, the SSN typically predicts fast dynamics in both high-variability and low-variability states (Supplementary Figure S2A). Even when the SSN shows some slowing at the lowest levels of input, due to the restoring-force couplings dipping below 1 (as in Figure 2E; the relaxation time in a direction with restoring coupling λ is *τ*/|λ| where τ is a cellular time constant), it transitions to fast dynamics (|λ|’s ≥ 1) for relatively weak input for which variability is still high relative to the high-input state (Supplementary Figure S2A) – a key distinction from the other models. Consistent with the SSN, in mouse V1, the decay of response back to spontaneous levels (or lower) after optogenetically-induced sudden stimulus offset is fast, occuring over 10 ms (Reinhold et al., 2015).

In summary, the SSN robustly captures multiple aspects of stimulus modulation of correlated variability and suggests a dynamical regime that uniquely captures a wide array of behaviors of sensory cortex.

## Experimental procedures

The values of all the parameters mentioned below are listed in Table 1.

**Table 1.**
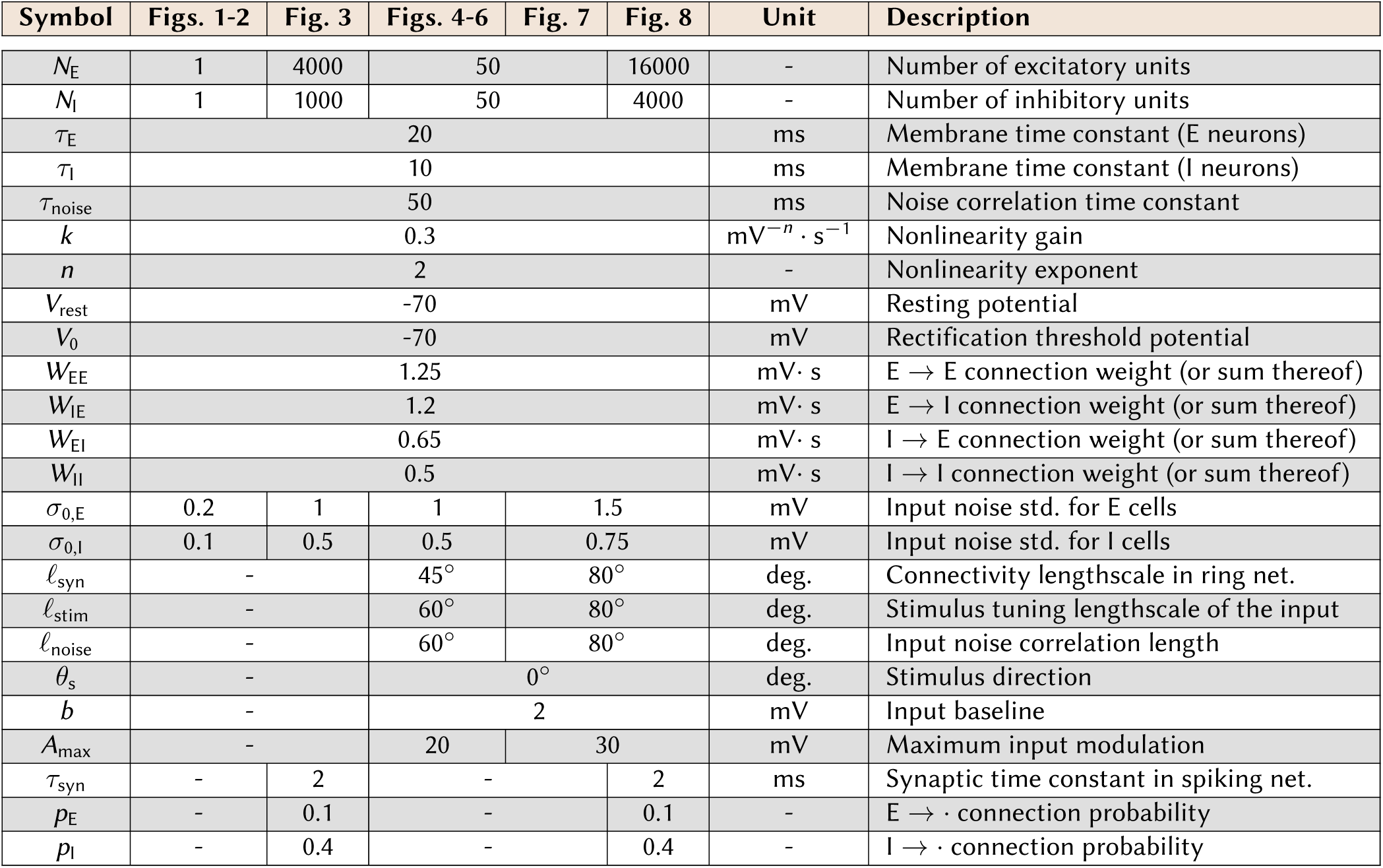
Parameters used in our simulations.

### Rate model

Our rate-based networks contained *N*_E_ excitatory and *N*_I_ inhibitory units, yielding a total *N*≡*N*_E_ + *N*_I_ The circuit dynamics were governed by Equation 1, which we rewrite here for convenience:

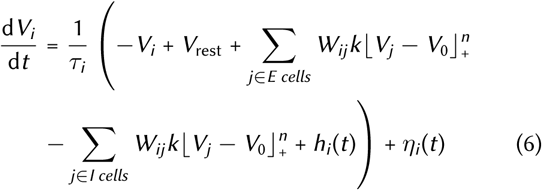

where *η_i_*(*t*) modelled fluctuations in external inputs (see below, “Input noise”). In all the figures of the main text, the exponent of the power-law nonlinearity was set to *n* = 2. The SI explores more general scenarios.

#### Mean external drive

In the reduced rate model of Figure 1, each unit received the same constant mean input *h*. In the ring model, the mean input to neuron i was the sum of two components,

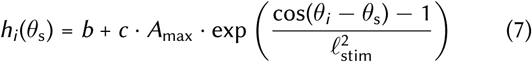

The first term *b* = 2 mV is a constant baseline which drove spontaneous activity. The second term modelled the presence of a stimulus moving in direction *θ*_s_ in the visual field as a circular-Gaussian input bump of width 𝓁_stim_ centered around *θ*_s_ and scaled by a factor *c* (increasing *c* represents increasing stimulus contrast), taking values from 0 to 1, times a maximum amplitude *A*_max_. We assumed for simplicity that E and I cells are driven equally strongly by the stimulus, though this could be relaxed.

#### Input noise

The input noise term *η*_i_(*t*) in Equation 6 was modelled as a multivariate Ornstein-Uhlenbeck process:

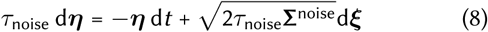

where d*ξ* is a collection of *N* independent Wiener processes and Σ^noise^ is an *N* × *N* input covariance matrix. Note that Equation 8 implies 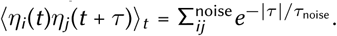

In the reduced model, noise terms were chosen uncorrelated,i. e. 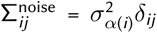. (where *δ*_*ij*_ = 1 if *i* = *j* and 0 otherwise), *α*(*i*) is the E/I type of neuron *i*, and 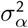 is the variance of noise fed to population *α* ∈ { E,I} (see Equation 10 below). In the ring model, the noise had spatial structure, with correlations among neurons that decreased with the difference in their preferred directions following a circular-Gaussian:

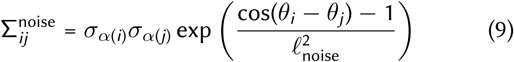

where *Θ_i_* and *Θ_j_* are the preferred directions of neurons *i* and *j* (be they exc. or inh.), and *𝓁*_noise_ is the correlation length (Table 1). The noise amplitude was given the natural scaling

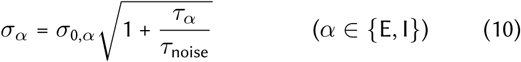

such that, in the absence of recurrent connectivity (**W** = 0), the input noise alone would have driven *V*_m_ fluctuations of standard deviation *σ*_0,E_ or *σ*_0,I_, measured in mV, in the E or I cells, respectively. We chose values of *σ*_0,*E*_ that yielded spontaneous Fano factors in the range 1.3-1.5 where appropriate, and chose 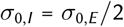 to make up for the difference in membrane time constants between E and I cells (Table 1).

#### Connectivity

The synaptic weight matrix in the reduced model was given by Equation 3 with synaptic strengths listed in Table 1. In the ring model, connectivity fell off with distance on the ring, following a circular-Gaussian profile:

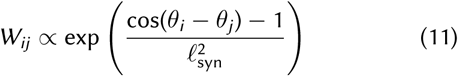

The connectivity matrix **W** was further rescaled in each row and in each quadrant, such that the sum of incoming E and I weights onto each E and I neuron (4 cases) matched the values of *W*_EE_, *W*_IE_, *W*_EI_ and *W*_II_ in the reduced model.

#### Simulated spike counts

To relate the firing rate model to spiking data (Figures 4, 6 and 7), we assumed action potentials to be emitted as inhomogeneous (doubly-stochastic) Poisson processes with time-varying rate 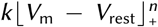. Spikes did not “re-enter” the dynamics of Equation 6, according to which neurons influence each other through their firing rates. Spikes were counted in 100 ms time bins and spike count statistics such as Fano factors and pairwise correlations were computed the standard way.

#### Theory of variability

To compute the moments of *V*_m_ analytically, we used i) a linear theory which assumes small fluctuations (the single-neuron gain function is Taylor-expanded to first order around the mean; Equation 4) and returns closed-form analytical results through standard multivariate Ornstein-Uhlenbeck theory (e.g. Renart et al. (2010); Tetzlaff et al. (2012); Hennequin et al. (2012); see SI for details), and ii) a nonlinear theory which does not rely on linearization, can handle large fluctuations and non-stationary transients, by assuming that variability in *V*_m_ is jointly Gaussian across neurons. We have used the nonlinear theory throughout the figures in this paper to smooth out the data points obtained numerically. The details will be published elsewhere (Hennequin and Lengyel, *in prep*.).

#### Mathematical definition of the “shear and restoring forces”

To uncover the structure of the forces acting on activity fluctuations, we focused on the linearized dynamics of Equation 4 and performed a Schur decomposition of the Jacobian matrix which included both the single-neuron leak and the effective connectivity (Murphy and Miller, 2009; Hennequin et al., 2012). In the reduced model, this amounted to expressing the dynamics of the E and I units in a different coordinate system, comprised of the two axes of E/I imbalance (thereafter called difference mode) and total activity (sum mode) depicted in Figure 2A–C. In that basis, the effective connectivity matrix – in which we also included the leak term – had a triangular (i.e. feedforward) structure. The diagonal contained the two eigenvalues of the effective connectivity matrix, and were interpreted as “restoring forces” due to their effect of pulling activity along each axis back to the mean. The upper triangular element of the Schur matrix was interpreted as a “shear force”, because it induced an effective connection from the difference mode onto the sum mode, resulting in the orange force field depicted in Figure 2B–C. We note that the Schur vectors are not pure, but instead weighted, sum and difference modes. Moreover, the elements of the Schur triangle are complex numbers in general; nevertheless, the intuition built in Figure 2 holds in the complex case too, because only the moduli of these complex numbers matter in computing total variability (in the limit of slow input noise). This is all detailed in the SI, together with explicit formulas for the input dependence of both shear and restoring forces, as well as how each force affects variability in the network. The SI also explains the higher-dimensional Schur decomposition performed on the effective ring connectivity (Figure 5), which is similar conceptually but demanded more involved treatment.

### Spiking model

#### Dynamics

In the spiking model, neuron *i* emitted spikes stochastically with an instantaneous probability equal to 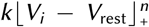, consistent with how (hypothetical) spikes were modelled in the rate-based case (cf. above). Presynaptic spikes were filtered by synaptic dynamics into exponentially decaying postsynaptic currents (E or I):

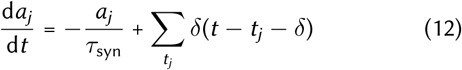

where the *t_j_*’s are the firing times of neuron *j*, *τ*_syn_ = 2 ms, and *δ* = 0.5 ms is a small axonal transmission delay (which also enables the distribution of the simulations onto multiple compute cores following Morrison et al., 2005, using custom software written in OCaml and linked to the MPI parallelization library). Synaptic currents then contributed to membrane potential dynamics according to

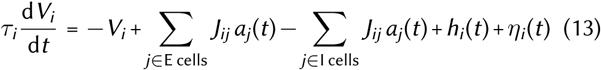

where the synaptic efficacies *J*_ij_ are described below, and the noise term *η*_i_ was modelled exactly as in the rate-based scenario. In Figure 3, the input noise covariance was simply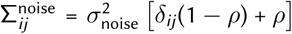. In Figure 8, input correlations were given again by Equation 9.

#### Connectivity

In Figure 3, for each neuron *i*, we drew *p*_E_*N*_E_ excitatory and *p*_I_*N*_I_ inhibitory presynaptic partners, uniformly at random. Connection densities were set to *p*_E_ = 0.1 and *p*_I_ = 0.4 respectively. The corresponding synaptic weights took on values 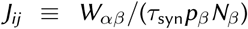 where 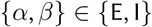 denote the populations to which neuron *i* and *j* belong respectively, and *W_αβ_* are the connections in the reduced model (Table 1). This choice was such that, for a given set of mean firing rates in the E and I populations, average E and I synaptic inputs to E and I cells match the corresponding recurrent inputs in the rate-based model. Synapses that were not drawn were obviously set to *J*_*ij*_ = 0.

To wire the spiking ring network of Figure 8, for each neuron i we also drew *p*_E_*N*_E_ excitatory and *p*_I_*N*_I_ inhibitory presynaptic partners, though no longer uniformly. Instead, we drew them from a (discrete) distribution over presynaptic index *j* given by:

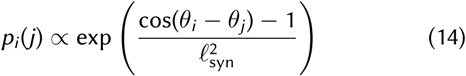

which mirrored the dependence of *W*_*ij*_ on angular distance in the rate model (cf. Equation 11). In Equation 14, “α” means this distribution is not normalized; we used simple box (rejection) sampling to draw from it. Synapses that were drawn took on the same values 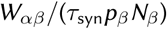 as in the randomly connected network (cf. above), again to achieve approximate correspondance with the rate model.

### Factor analysis

We performed factor analysis of spike counts, normalized by the square root of the mean spike count for each neuron. This normalization was such that the diagonal of the spike count covariance matrix **C** contained all the single-neuron Fano factors, which is the usual measure of variability in spike counts. In the ring model, such a normalization also prevented **C** from being contaminated by a rank-1 pattern of network covariance merely reflecting the tuning of single-neuron firing rates (the “Poisson” part of variability, which indeed scales with the mean count), but instead expressed covariability in the above-Poisson part of variability in pairs of cells. Factor analysis decomposes **C** as **C**_private_ + **C**_shared_, where **C**_shared_ has much lower rank than **C**_private_. Here, since we could simulate to model long enough to get a very good estimate of the spike count covariance matrix **C**, we performed factor analysis by direct eigendecomposition of **C**, thus defining 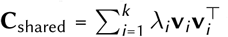 whereby the top *k* eigen-vectors **v**_1_,…., **v**_k_ of **C** contributed to shared variability in proportion of the corresponding eigenvalues λ_i_. We kept *k* = 1 eigenmode for the two-population model of Figure 3, as we found the first eigenvalue of **C** to be singled out (much larger than all other eigenvalues) across all values of *ρ* and *h*. For the ring model of Figure 8, between 3 (for large *c*) and 5 (for small *c*) eigenvalues of **C** stood out. We chose to keep *k* = 5 modes in order to conservatively estimate the drop in shared variability.

## Acknowledgments

This work was supported by NIH grant R01-EY11001 (K.D.M.), the Gatsby Charitable Foundation (K.D.M.), Medical Scientist Training Program grant 5 T32 GM007367-36 (D.B.R.), the Swartz Program in Computational Neuroscience at Columbia University (Y.A.), the Postdoc Program of École des Neurosciences, Paris, France (Y.A.), and the Wellcome Trust (M.L.,G.H.). We thank Larry Abbott and Andrew Tan for helpful discussions. Y.A. would like to thank David Hansel and the Centre de Neurophysique, Physiologie, et Pathologie, Paris, for their hospitality.

## S1 Recap of model setup

We consider the stochastic and nonlinear rate model of Equation 1 of the main text. To simplify notations, we assume *V*_rest_ = 0 mV without loss of generality as it can be absorbed in the external input, and rewrite:

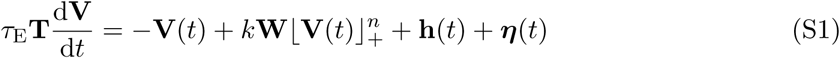

with *n* > 1 (*n* = 2 throughout the main text). In Equation (S1), 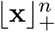 denotes the pointwise application of the threshold power-law nonlinearity to the vector x, that is, 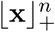 is the vector whose *i^th^* element is 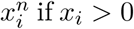 0, or 0 otherwise; **T** is a diagonal matrix of relative membrane time constants measured in units of *τ*_E_; **W** is a matrix of synaptic connections, made of *N*_E_ positive columns (corresponding to excitatory presynaptic neurons) and Ni negative columns (inhibitory neurons) for a total size of *N* = *N*_E_ + *N*_I_; **h**(*t*) is a possibly time-varying but deterministic external input to neuron *i*; and ***η*** is a multivariate Ornstein-Uhlenbeck process with separable spatiotemporal correlations given by

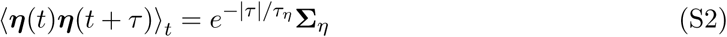

where Σ_*η*_ is the covariance matrix of the input noise and *τ_η_* is its correlation time. In particular, we are going to study how *τ_η_* and correlations in Σ_*η*_ affect network variability. We adopt the following notations for relative time constants:

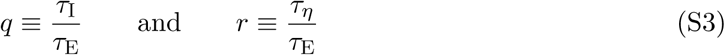

In general, recurrent processing in the network is prone to instabilities due to the expansive, non-saturating *V*_m_-rate relationship in single neurons. However, there are generous portions of parameter space in which inhibition dynamically stabilizes the network. We refer to this case as the “supralinear stabilized network”, or SSN (Ahmadian et al., 2013; Rubin et al., 2015).

## S2 Mean responses in the stabilized supralinear regime

### S2.1 Recap of Ahmadian et al. (2013)’s theoretical analysis

Our analysis of the stochastic SSN developed in Section S3 will show that the modulation of variability relies on the nonlinear behavior of *mean* responses to varying inputs (Figure 1D of the main text), which were studied previously (Ahmadian et al., 2013). In particular, the transition from superlinear integration of small inputs to sublinear responses to larger inputs (Figure 1 of the main text) could be explained using simple scaling arguments, which we briefly reproduce here. Note that here we have written the circuit dynamics in voltage form (Equation (S1)), while Ahmadian et al., 2013 chose a slightly different rate form; accordingly, the equations we now derive differ from the original equations in their form, but not in their nature (in fact steady state solutions studied in Ahmadian et al., 2013 are mathematically equivalent in the two formulations, and moreover when **T** is proportional to the identity matrix, dynamic solutions are also exactly equivalent (Miller and Fumarola, 2011)).

This section is devoted to mean responses, therefore we neglect the input noise ***η*** for now. We thus write the deterministic dynamics of the mean potentials *V̅*_*i*_ as

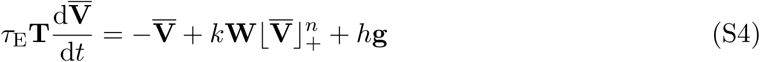

and ask how neurons collectively respond to a constant external stimulus *h* fed to them through a vector g ~ 𝒪( 1) of feedforward weights. Perhaps after some transient, and assuming the network is stable (see below), the network settles in a steady state **V̅** which must obey the following fixed point equation, obtained by setting the l.h.s. of Equation (S4) to zero:

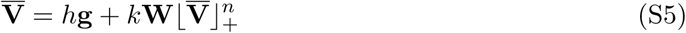

As in the main text, we focus on the case of a threshold-quadratic nonlinearity, *n* = 2, though the following derivations can be extended to arbitrary *n* > 1. Following Ahmadian et al. (2013), we begin by writing **W** ≡ Ψ *φ***J** where 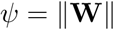 for some matrix norm 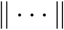, and the dimensionless vector **J** has 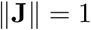. We also define dimensionless mean voltage and input respectively as

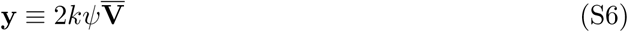

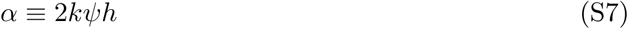

(note that the definition of *α* differs from that in Ahmadian et al., 2013 by a factor of 2). With these definitions, the fixed point equation for the mean potentials, Equation (S5), becomes

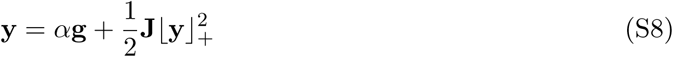

**Network responses to small inputs** When *α* is small (i.e. *h* is small, given fixed connectivity strength *Ψ*), it is easy to see that

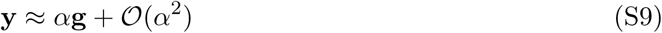

In essence, the fixed point Equation (S8) is already the first-order Taylor expansion of y for small *α* (indeed, the recurrent term 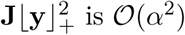, self-consistently). Thus, for small input *α,* membrane potentials scale linearly with *α,* and firing rates are quadratic in *α,* merely reflecting the single-neuron nonlinearity. In other words, the network behaves mostly as a relay of its feedforward inputs, with only minor corrections due to recurrent interactions.

More generally, by repeatedly substituting the right side of Eq. S8 for y in Eq. Eq. S8, we arrive at the expansion

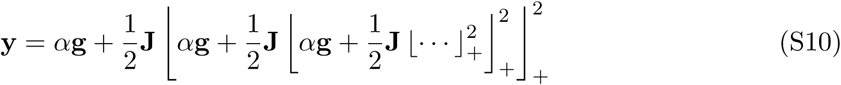

The net result involves a series of terms of order *α*, *α*^2^, *α*^4^ …, which can be expected to converge for small *α* (*α*≪ 1).

**Network responses to larger inputs For large** *α* (*α*≫ 1), the expansion of Eq. S10 will not converge and so cannot describe responses. Physically this tends to correspond to the excitatory subnetwork becoming unstable by itself. At the level of the fixed point equation S8, recurrent processing involves squaring **V̅**, passing it through the recurrent connectivity, adding the feedforward input, squaring the result again, …, which for large enough input and purely excitatory connectivity would yield activity that grows arbitrarily large. A finite-activity solution is achieved through stabilization by inhibitory feedback. Mathematically, for this to occur, the recurrent term 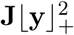 must cancel the linear dependence of y on *α* in Eq. S8 (since any linear dependence would be squared by the right side of Eq. S8, then squared again, …, to yield an explosive series like Eq. S10). That is, we must have

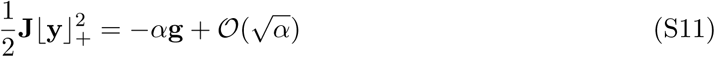

such that (again from Equation (S8))

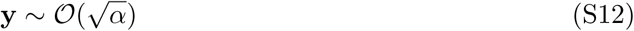

at most. This means that membrane potentials scale at most as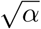, i.e. firing rates scale at most linearly in *α*. However, in many cases, firing rates too will be sublinear in *α*. This is best examplified in the context of our two-population E/I model, by following Ahmadian et al. (2013) and introducing the notation:

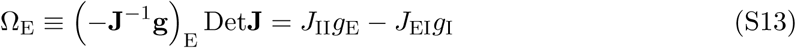

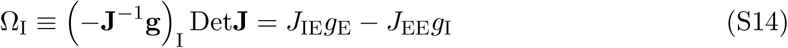

(note that we only consider networks in which Det**J** > 0, as it must for stabilization to occur for all input levels *α*, Ahmadian et al. (2013)). Equation (S11) can then be rewritten as

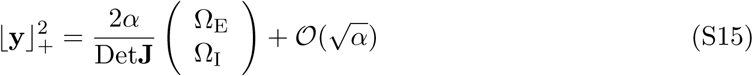

Now, depending on the choice of parameters (recurrent weights **J** and feedforward weights g), Ω_E_ in particular can be negative. Since 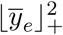, is positive, it must be that the sublinear term 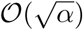, dominates over the (negative) linear term 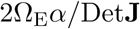, at least over some range of *α* over which the E firing rate is non-zero. In this case, 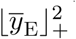, behaves roughly as 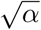 over some range^1^ before it gets pushed to zero, and accordingly *y̅*_E_ must be approximately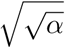 over the same range, i.e. the E unit responds strongly sublinearly. Ahmadian et al. (2013) referred to this regime of eventual decrease of *y̅*_E_ with increasing stimulus strength as “supersaturation”, and showed that it occurs for physiologically plausible parameter regimes. Our choice of parameters for the two-population model of the main text falls within this class of strongly sublinear E responses (Ω_E_ < 0), but we will show in Section S3 that the SSN displays the same input modulation of variability irrespective of the sign of Ω_E_.

In summary, the SSN responds superlinearly to small inputs, and sublinearly to larger inputs. Firing rates become at most linear (but will be sublinear if Ω_E_ < 0) with large inputs. Accordingly, membrane potentials show a transition from linear to (potentially strongly) sublinear responses to increasing inputs. Moreover, this transition occurs for *α* ∼ 𝒪 (1).

### S2.2 What do we expect for typical networks?

In the context of the reduced two-population model of the main text, we now complement the above theoretical arguments with a numerical analysis of the SSN’s responses across a wide range of parameters, in order to form a picture of the “typical” behavior of the SSN in physiologically realistic regimes. We will later (Section S3) reuse these numerical explorations to show that the modulation of variability by external input in the SSN is robust to changes of parameters.

The dynamics of the trial-averaged dimensionless “population voltages” are given by

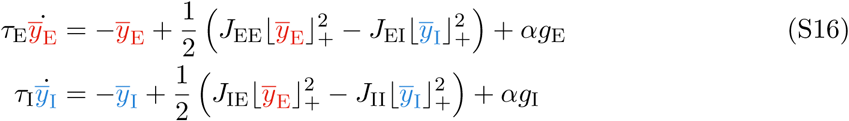

It is difficult to get good estimates of the values of the 6 free parameters (feedforward weights and recurrent weights) directly from biology. Therefore, our approach is to construct a large number of networks by randomly sampling these parameters within broad intervals, and rejecting those networks that produce unphysiological responses according to conservative criteria that we detail below. We then examine the behavior of each of these networks and perform statistics on the various kinds of responses that have been identified in the theoretical analysis of Section S2.1.

We thus constructed 1000 networks by sampling both feedforward weights {*g_α_*} and recurrent weights {*J_αβ_*} (for *α,β* ∈ {E,I}) uniformly from the interval [0.1; 1], and subsequently normalizing their (vector) L_∞_-norm such that max(*g_α_*) = max(*J_αβ_*) = 1. We then sampled the overall connectivity strength *ψ* (cf. Section S2.1) from the interval [0.1; 10]. This interval was based on rough estimates of the average number of input connections from the local network per neuron (between 200 and 1000), average PSP amplitude (between 0.1 mV and 0.5 mV) and decay time constants (5 to 20 ms), giving a range of connectivity strengths – which in our model is the product of these three quantities – between 0.1 and 10 mV/Hz.

Instead of choosing a range of *α* and simulating the dynamics of Equation (S16) to compute mean voltages, we instead observed that increases monotonically with *α* and for each network we chose a range of *y̅*_I_ corresponding to mean I firing rates ((*y̅*_I_/2*Ψ*)^2^/*k*) in the range [0; 200] Hz, thus assuming that mean I responses above 200 Hz would be unphysiological. For each *y̅*_I_ in this discretized range we solved for *y̅*_E_ analytically by noting that the input *α* can be eliminated from the pair of fixed-point equations (Equation (S16) with l.h.s. set to zero), yielding a fixed-point curve in the (*y̅*_E_,*y̅*_I_) plane:

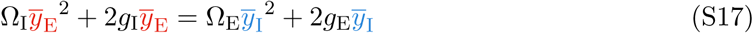

Given *y̅*_I_ it is easy to solve this quadratic equation for *y̅*_E_. We rejected those parameters sets for which we encountered either i) complex solutions for *y̅*_E_, or ii) real but unstable solutions, as assessed by the stability conditions Tr*𝒥* < 0 and Det *𝒥* > 0.01 (with the Jacobian matrix *𝒥* defined in Equations (S19) and (S21)), or iii) stable solutions that involved E firing rates ((*y̅*_E_/2*Ψ*)^2^/k) either greater than 200 Hz, or smaller than 1 Hz for the largest value of *y̅*_E_. Finally, for each fixed point (*y̅*_E_,*y̅*_I_), we computed the corresponding *α* from either of the two fixed-point equations (Equation (S16) with l.h.s. set to zero), e.g. 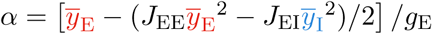. This procedure was numerically much more efficient than simulating the dynamics of Equation (S16) until convergence to steady-state.

The parameters of the retained networks spanned a large chunk of the invervals in which they were sampled (Figure S1A and B). Because stability for large *α* requires Det**J** > 0, i.e. *J_EI_J_IE_* > *J_EE_J_ii_*, the largest of all sampled *J_αβ_*’s was often either *J_EI_* or *J_IE_* which then, due to the *L*_∞_-norm normalization, assumed a value of one (Figure S1A). We also observed that the input weight *g_E_* was often larger than *g_I_* (Figure S1B). About 90% of the sampled networks has Ω_E_ > 0, implying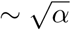 scaling of *y̅*_E_ and *y̅*_I_ for large *α* (example in Figure S1D, top). In these networks, E and I rates were linear in *α* for *α* large enough, and so were also linear in each other when large enough (Figure S1E, black). The rest of the networks (10%) had Ω_E_< 0 and therefore showed supersaturation of the E firing rate for large input (Figure S1D, bottom) and E responses that were sublinear in I responses (Figure S1E, orange).

It is worth noting that for networks with small overall connectivity strength *ψ*, the proportion of Ω_E_ < 0 and Ω_E_ > 0 cases tend to even out (Figure S1C). This is because, for supersaturating networks, the peak E firing rate is inversely proportional to *ψ*^2^ (Ahmadian et al., 2013), so for large *ψ* the peak firing rate is low and therefore the final value of *r̅*_E_ reached for *r̅*_I_ = 200 Hz likely falls below our threshold of 1 Hz, resulting in a rejection of the parameter set.

In sum, the nonlinear properties of the SSN’s responses to growing inputs, summarized in Section S2.1, are robust to changes in parameters so long as these keep the network in a regime “not too unphysiological” in a conservative sense. Using the same collection of sampled networks, we will show below that the modulation of variability with input described in the main text is equally robust to parameter changes.

## S3 Activity variability in the two-population SSN model

In this section, we derive the theoretical results regarding activity variability in the two-population model of the main text. We use these analytical results to demonstrate robustness of our results to changes in parameters, which we also verify numerically using the collection of networks with randomly sampled parameters introduced in Section S2.2.

### S3.1 Linearization of the dynamics

We now consider the noisy dynamics of the two-population model of the main text in which the E and I units represent the average activity of large E and I populations. To study variability analytically, we linearize Equation (S1) around the mean, thus examining the local behavior of small fluctuations *δ***V**:

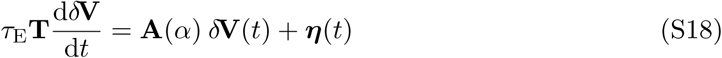

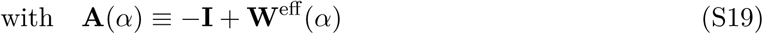

The effective connectivity **W**^eff^ depends on the (dimensionless) input *α* through its dependence on mean responses, following

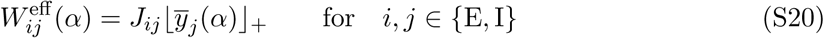

where we have used the definition of the dimensionless voltage y and dimensionless connections **J** introduced in Section S2.1. With our notations, the Jacobian matrix

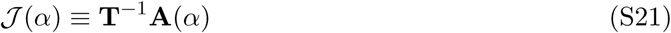

is unitless, so that, e.g., the interpretation of a real negative eigenvalue λ of *𝒥* is that the corresponding eigenmode decays asymptotically with time constant *τ*_E_/|λ| as a result of the recurrent dynamics. We parameterize the input noise covariance as

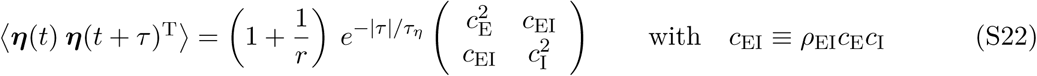

such that, in the limit of small *α* – in which the network is effectively unconnected, because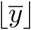 in Equation (S20) is small – the E unit has variance 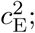 the I unit then has variance 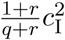. The parameter *ρ*_EI_ determines the correlation between input noise to the E and I units.

### S3.2 General result

As shown in the appendix, the full output covariance matrix 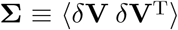 can be calculated by solving a set of linear equations, which yields:

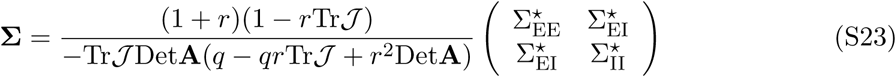

with

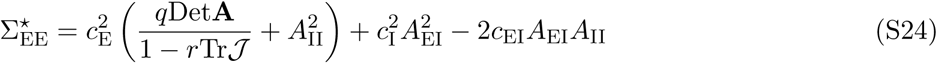

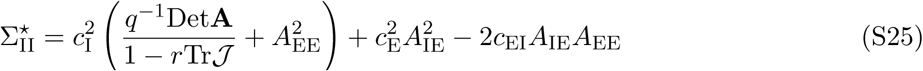

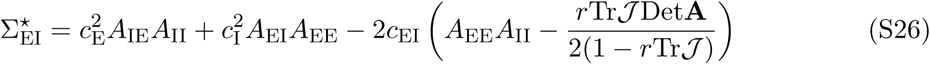

In Equations (S23) to (S26), each term that depends on **A** or *𝒥* depends implicitly on the (dimensionless) constant input *α* delivered to both E and I populations, because **A** (or *𝒥*) depends on mean voltages (through Equation (S20)) which themselves depend on *α*. Note also that, for the network to be stable at a given input level *α*, the Jacobian matrix *𝒥*(*α*) should obey Tr*𝒥* < 0 and Det*𝒥* > 0 (with the latter equivalent to Det**A** > 0).

Among other things, we will analyze the behaviour of the total variance, i.e. the trace of Σ given by

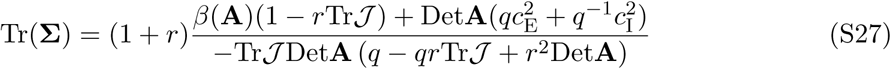

with **A** defined in Equation (S19) and

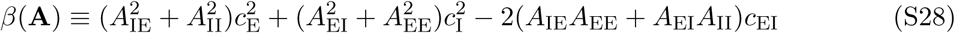

### S3.3 Analysis in simplified scenarios

In order to understand what Equation (S27) tells us about the modulation of variability with the input *α*, we make a couple of assumptions that greatly simplify the expression for the total variance with little loss of generality. First, we consider the limit of slow^2^ input noise which we find empirically is approached rather fast, with *τ_η_* = 50 ms already giving a close approximation given *τ*_E_ = 20 ms and *τ*_E_ = 10 ms. Next, we assume that

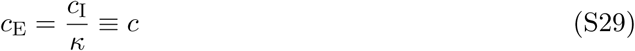

and *ρ*_EI_ = 0, i.e. the E and I units have uncorrelated input fluctuations of equal amplitude (the impact of positive input correlations, *ρ*_EI_ > 0, will be discussed in Section S3.4). With these two assumptions, the total variance simplifies into

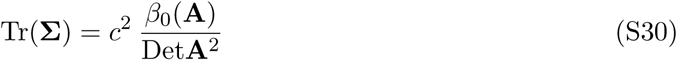

which provides a good basis for discussion. Here we defined *C*^2^ *β*_o_(**A**) to be *β*(**A**) with *c*_EI_ set to zero. The typical behavior of 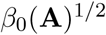 and Det**A** is shown in Figure S2A. Both can be expressed as a function of mean responses using Equations (S19) and (S20):

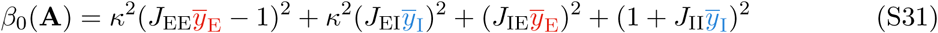

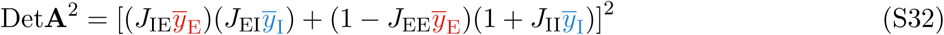

Note that to simplify notations we have dropped the 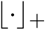 that should surround every *y̅*. Based on these expressions, we now examine the behavior of variability in the small and large *α* limits and show that the total variance should typically grow and then decay with increasing *α,* and therefore should exhibit a maximum which empirically we find occurs for *α* ∽ 1.

#### Behavior of the total variance for small *α*

Using Equations (S30) to (S32), we find the slope of the total variance at *α* = 0 to be

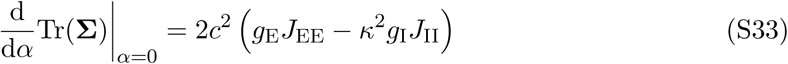

Thus, when the noise power fed to inhibitory cells is sufficiently small, 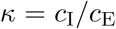will be small enough that the expression in Equation (S33) will stay positive, and therefore total variability will grow with small increasing *α*. Indeed, we find that this happens for most (> 90%) of the randomly sampled networks of Section S2.2 with *κ* as large as 1/2 (Figure S2A, bottom). Moreover, restricting the analysis to the E unit gives 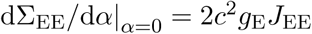 which is always positive, independently of *k*. Thus, for slow enough input noise, the variability in the E unit always increases with small *α*.

We can extend this argument to slightly larger values of *α* by further inspecting the numerator and denominator in Equation (S30). Although the first term in the numerator, (*J*_EE_*y̅* − 1)^2^, originally decays with *α* as *y̅*_E_ grows from 0 to 1/*J*_EE_, the other three terms always grow with *α* as long as mean voltages do, and thus we expect the numerator to typically grow. This is indeed what we find in all sampled networks (Figure S2A). On the other hand, the denominator (Equation (S32)) is the square of the sum of two terms, the first one initially small and growing, and the second one initially large and decaying. Indeed, the second term starts at 1 for *α* = 0, because the *y̅* terms are all zero, and then decays to zero as the network enters the inhibition-stabilized (ISN) regime and the effective excitatory feedback gain *J*_EE_*y̅*_E_ becomes larger than one^3^ (Tsodyks et al., 1997; Ozeki et al., 2009). Thus, due to this partial cancellation of growing and decaying terms, we expect the denominator to either decrease, or grow very slowly, with increasing *α* (Figure S2A), until it starts growing faster (see arguments below for the large *α* case) in the very rough neighborhood of the ISN transition. All in all, the ratio of a fast growing numerator to a slower growing denominator suggests that the total variance should robustly grow with small increasing *α* (Figure S2A, bottom).

#### Behavior of the total variance for large *α*

As the input grows, so do the mean (dimensionless) voltages *y̅*_E_ and *y̅*_I_ at least over some range of *α*. Therefore, we expect *both* the numerator *and* the denominator that make up the total variance in Equation (S30) to grow with large enough and increasing *α*. However, loosely speaking, the numerator grows as *y̅* ^2^ while the denominator grows as *y̅* ^4^, which can be seen by inspecting Equations (S31) and (S32). Thus, their ratio should decrease roughly as 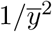.

This argument can be made more rigorous in the case Ω_E_ > 0, i.e. when the E unit does not supersaturate. In this case, from Equation (S15) we have 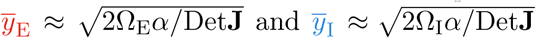 for *α* large enough. Therefore, in the large *α* limit, the numerator and denominator of Equation (S30) behave as

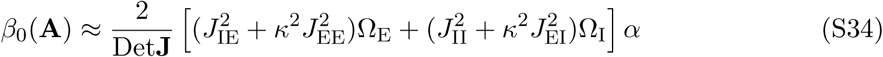

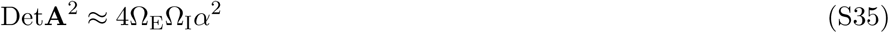

respectively, therefore the total variance (their ratio) decreases as 1/*α*. For Ω_E_ <0, the large *α* limit is irrelevant strictly speaking, as in this limit 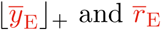 go to zero. In this case the total variance does not decrease asymptotically but reaches a finite limit of 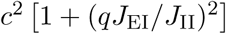. However, we find empirically that the peak of variability always occurs well before the onset of supersaturation, in a regime where both *y̅*_E_ and *y̅*_I_ are still growing with *α* while remaining roughly proportional to each other (Figure S1E), so that the argument made above can be repeated: the total variance decreases as 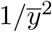 for a while after having peaked.

#### Where does variability peak?

The above arguments, derived for slow noise *τ_η_* → ∞, show that growing inputs typically increase, and then suppress, total variability in the two-population SSN. Thus, total variability (and even more certainly, variability in the E unit) typically exhibits a maximum for some intermediate value of *α*. We find empirically that, even for finite *τ*_*η*_, the location of this variance peak is well approximated by its location in the limit of fast inhibition, *q* → 0, which we can estimate analytically. Indeed, in this limit, the I cell responds instantaneously to changes in E activity and input noise, such that

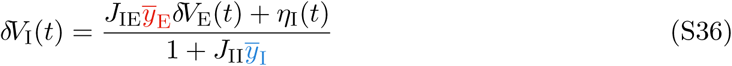

Consequently, *δV*_E_ now obeys one-dimensional dynamics given by

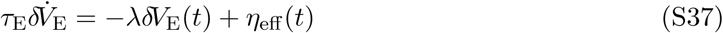

where

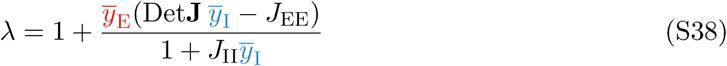

and *η*_eff_ is a noise process (a linear combination of *η*_E_ and *η*_I_) with temporal correlation length *τ_η_* and a variance that is empirically irrelevant for the arguments below^4^ In this case, the variance of *δV*_E_ is inversely proportional to 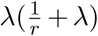, and therefore should be maximum at the input level *α* that minimizes λ. Observing from Figure S1E that *y̅*_E_ and *y̅*_E_ are roughly proportional over a large range of *α* (for Ω_E_ < 0), if not the entire range (for Ω_E_ > 0), we can make the following approximation:

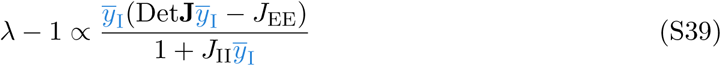

whose minimum is straightforward to calculate and is attained for

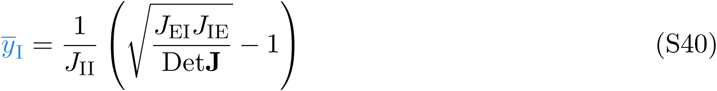

We find that the *α* of maximum variance in the E unit is indeed very well approximated by the *α* at which *y̅*_I_ reaches the threshold value of Equation (S40), especially in the absence of input correlations (*ρ*_EI_ = 0, Figure S2B, left). For correlated noisy inputs, the criterion of Equation (S40) deteriorates slightly but still consistently provides an upper bound on the *α* of maximum E variance (Figure S2B, right).

Interestingly, the criterion for maximum variance in Equation (S40) is equivalent to a criterion about the effective I→I connection, given by 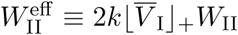 (cf. main text Equation (6)). Specifically, at the peak of variance we expect to have

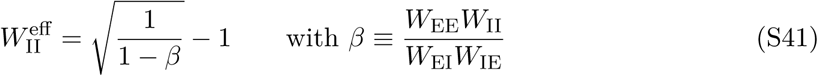

where *β* < 1 is in some sense the ratio of what contributes positively to the activity of the E cell (product of self-excitation *W*_EE_ with disinhibition *W*_II_) to what contributes negatively to it (the product *W*_IE_*W*_EI_ quantifying the strength of the E → I → E inhibitory feedback loop). Thus, in networks with inhibition-dominated connectivity, i.e. ones in which *β* ≪ 1, we expect 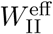 to reach the criterion of Equation (S41) earlier as the input grows (this argument implictly assumes that the rate of growth of 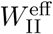 itself doesn’t depend too much on *β*, which we could confirm numerically).

Finally, we note that since variability peaks for *α* ∽ 𝒪(1) and *y* ∽ 𝒪(1), networks with stronger connectivity (large *ψ*) will exhibit a peak of variance for smaller external input *h* (because *α* ã *ψh*) – and this peak will occur for lower voltages/firing rates (because *V̅ αy*/*ψ*).

### S3.4 Effects of input correlations

To see the effect of input correlations on variability, we return to the expression for Σ_EE_ in Equation (S27), assume again that *τ_η_* →∞ and 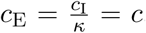, but now with *ρ*_EI_≠ 0. We thus obtain:

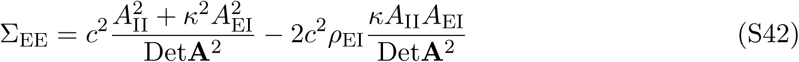

Thus, total E variability is equal to that without input correlation (the first term), minus a positive term proportional to *ρ*_EI_. Thus, positive input correlations always decrease variability in the E unit (and, in particular, its peak; Figure S2C, right), while negative correlations increase it. Moreover, the subtracted term has the same large-α behavior as the first term, because the two terms share the same denominator and for large alpha both numerators are 𝒪(*y̅*_j_ ^2^). Thus, input correlations should not affect the qualitative, decreasing behaviour of E variance for large increasing inputs. For small *α* and large *ρ*_EI_, however, we expect 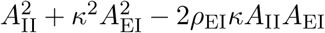 to grow much more slowly than 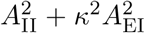; and indeed, in the extreme case *ρ*_EI_ = 1, the total numerator becomes 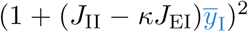, which can even decrease transiently with increasing *α* if *kJ*_EI_ >*J*_II_ (this occurs in about half of our thousand networks). This, in effect, shifts the peak of E variability to smaller values of *α* (Figure S2C, left).

The situation for the I unit is a bit different, as input correlations affect the I variance differently depending on whether the network has already made the transition to the ISN regime. Indeed, under the same assumptions as above, the I variance is given by

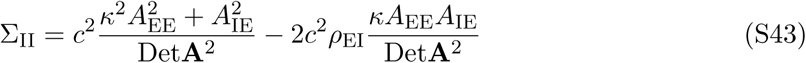

In the ISN regime, *A*_EE_ > 0, so that input correlations decrease I variability, just as it does for E variability as seen above. For small enough inputs, however, the network is not yet an ISN (*A*_EE_< 0), so that the effect of correlations is reversed: larger input correlations increase I variability.

In sum, input correlations modify the fine details of how large the variance grows and how early it peaks with increasing inputs, but they do not modify the qualitative aspects – in particular, the non-monotonic behavior – of variability modulation with external inputs in this two-population SSN model.

### S3.5 Mechanistic aspects: Schur decomposition

We now unpack the mechanistic aspects of variability modulation described in the main text, i.e. give mathematically precise meaning to the “forces” of Figure 2 (main text) acting on input fluctuations. We do this through a Schur decomposition (see e.g. Murphy and Miller, 2009 and its supplementary material in particular) of the 2-population model’s Jacobian matrix in Equation (S21):

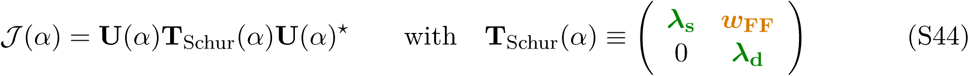

where ^.★^ denotes the conjugate transpose, λ_s_ and λ_d_ are the two (either real or complex-conjugate^5^) eigenvalues of *𝒥*(*α*), and the columns of **U** are the (orthonormal) Schur vectors such that **UU**^★^ = **U**^★^x2605;**U** = I. Expressing the E and I voltage fluctuations in the Schur basis as **z** ≡ **U**^★^ δ**V**, their dynamics become

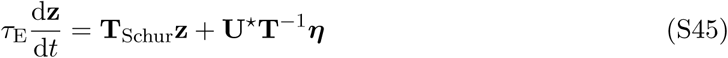

In the case of the 2-population E/I architecture considered here (**W** given by Equation 4 of the main text), the first Schur vector is a “sum mode” in the generalized sense (Murphy and Miller, 2009), i.e. its excitatory and inhibitory components have the same sign^6^.This corresponds to patterns of network activity in which the excitatory and inhibitory units are simultaneously either more active or less active than average. The second Schur mode is a generalized “difference mode” in that its excitatory and inhibitory components have opposive signs. (Hence the notations λ_s_ and λ_d_.) In theory, **U** depends on the input *α*, because *𝒥* does. However, we find that passed a relatively small value of *α*, the Schur vectors do not change much and are indeed sum-like and difference-like across all thousand networks studied in Sections S2 and S3 (Figure S2E).

The Schur decomposition reveals through **T**_Schur_(*α*) a feedforward structure hidden in the effective, recurrent connectivity *𝒥*(*α*): the difference mode feeds the sum mode with an effective feedforward weight *ω*_FF_ (also a complex number if the eigenvalues have an imaginary component), given by the upper right element of the triangular matrix **T**_Schur_. On top of this, both patterns inhibit themselves with the corresponding negative weight λ_d_ or λ_s_. Note that the sum of squared moduli (squared Frobenius norm 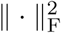) is preserved by the unitary transformation 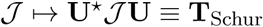, such that 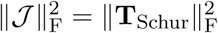, i.e.

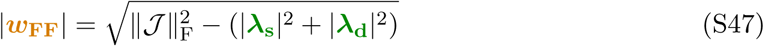

In the main text, we called the effect of λ_s_ and λ_d_ “restoring forces”, and that of *ω*_FF_ a “shear force”, because of the way they contribute to the flow of dynamics in the E/I activity plane and thus distort the ellipse of input fluctuations. Fluctuations are quenched along both the sum and the difference axes, in proportion of λ_s_ and λ_d_ respectively, and fluctuations along the difference axis are amplified along the sum axis in proportion of *w*_FF_.

The calculation of the network covariance matrix (Equation (S27)) can also be performed in the Schur basis, and doing this sheds further light on the roles of λ_d_, λ_s_ and *w*_FF_ in shaping variability. We begin by observing that

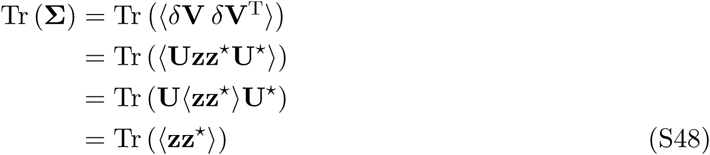

(the last step following from **UU**^★^ = **I**). Thus, the total variance is preserved in the Schur basis. Next, taking the Fourier transform of Equation (S45) and rearranging term yields

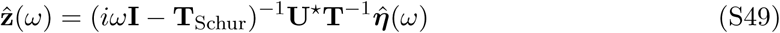

where 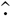 denotes the Fourier transform and 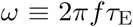 is a dimensionless frequency. Moreover, according to Parseval’s theorem we have

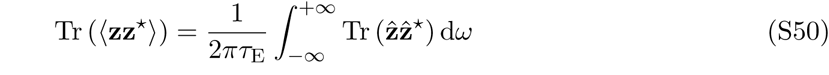

Thus, combining Equations (S48) to (S50) we get

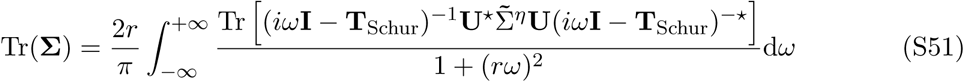

where 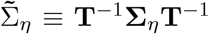. To simplify the calculation we now assume uncorrelated input noise terms, with the power of noise input to *E* and *I* balanced such that *κ* = *q* and 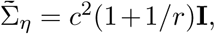, leading to:

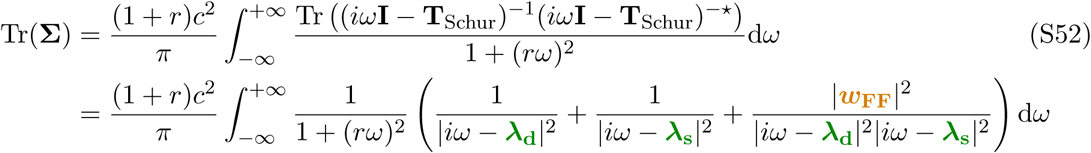

where the second equality comes from having inverted the upper-triangular matrix *iω* **I**—**T**_schur_ analytically and taken its squared Frobenius norm. Carrying out the integral gives

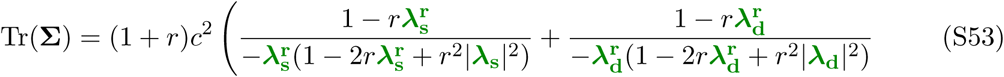

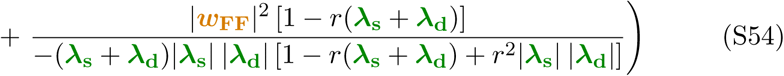

where 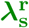 and 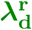 stand for the real parts of λ_s_ and λ_d_ respectively (they must both be negative for the dynamics to be stable).

This expression simplifies in the slow noise limit, *r* → ∞:^7^

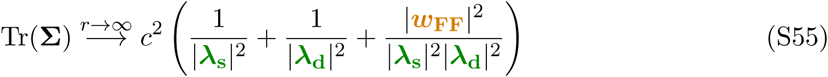

In this limit, the picture of the forces drawn in a plane of sum and difference activity (Figure 2 of the main text), assuming that they are real quantities, becomes accurate even when the eigenvalues of *𝒥* are complex-conjugate (in which case, as mentioned above in passing, the sum-like mode feeds back onto the difference mode, although this interaction is much weaker than the opposite one). Indeed, in Equation (S55), the elements of **T**_Schur_ are reduced to their moduli, so even when they are complex one can still interpret Equation (S55) as the total variance in a system with the same real Schur vectors, real eigenvalues equal to 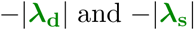 respectively, and a real feedforward weight equal to 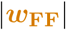.

Equation (S55) shows in more details how the shear and restoring forces contribute to variability. In loose terms, the total variance is a sum of two contributions: one that does not depend on *ω*_FF_ and decreases with 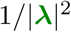, and one that grows with 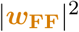 but is also divided by a term of order λ^4^ (where λ is a loose notation to denote the overall magnitude of the eigenvalues). Thus, as the input grows, the effect of the eigenvalues on variability becomes much stronger than that of balanced amplification. Such a dominance can also be understood from the structure of the force fields that negative self-couplings and balanced amplification induce. Restoring forces are proportional to the distance from the origin: the stronger the momentary *V*_m_ deviation from mean in any direction, the stronger the pull towards the origin in the same direction (main text Figure 2C, green arrows). In contrast, shear forces grow along the difference axis while pointing in the orthogonal, sum direction, such that larger deviations in the sum do not imply larger shear force (main text Figure 2C, orange arrows). Thus, self-inhibition leads to exponential temporal decay of activity fluctuations, whereas balanced amplification gives only linear growth. This explains why, for large enough input, *V*_m_ variability decreases with increasing input even when all forces grow in magnitude at the same rate (Figure S2A).

Equation (S55) also shows that if one of the eigenvalues transiently weakens with increasing input, then variability should transiently grow. This explains a large part of the variability peak observed in the network of the main text, and indeed, it also predicts variability growth in most of the thousand networks investigated here. However, there are cases where variability transiently grows, without any weakening of eigenvalues (Figure S3A). In those cases, setting *ω*_FF_ to 0 in Equation (S55) wrongly predicts purely decaying variability (compare dashed and solid black lines in Figure S3A, bottom). Thus, in general, initial variability growth results from the combined effects of weaker inhibitory self-couplings *and* strong balanced amplification.

### S3.6 How do the “forces” depend on the input?

The input dependence of the shear (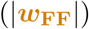) and restoring (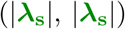) forces can be understood from the input dependence of mean responses (*y̅*_E_ and *y̅*_I_), which were examined previously in Section S2. First, at *α* = 0 (no input) the effective connectivity is zero, thus *𝒥* = diag(−1, _ *q*^−1^) and therefore the two eigenvalues are − 1 and _ 1/*q*. To see how the eigenvalues change with the input, let us note that for a 2 × 2 matrix, the sum of the eigenvalues is equal to the trace of the matrix while their product is equal to its determinant. Thus, when both eigenvalues are real (which they are for small enough *α*), both the arithmetic and geometric mean of 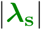 and 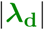 can be related to the elements of *𝒥*, which themselves depend directly on *y̅*_E_ and *y̅*_I_. This yields:

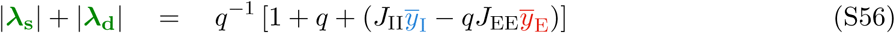

and

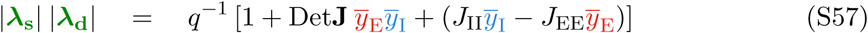

We see that, by both measures, the overall restoring force tends to grow with increasing input *α*, because i) mean responses grow too, and therefore so does the product term in Equation (S57), and ii) *y̅*_I_ tends to grow larger than *y̅*_E_ (Figure S1E), so that the weighted difference terms inside round brackets in both Equations (S56) and (S57) increase, at least for large enough *α*. However, when *g*_E_*J*_EE_ > *g*_I_*J*_II_, the difference term in Equation (S57) will initially grow negative with increasing – but small – *α*, before it increases again for larger *α*. This means that at least one of the eigenvalues will decrease. In such a case, whether or not *both* eigenvalues decrease transiently depends on the behavior of the difference term in Equation (S56). The requirement for this difference term to decrease initially is *qg*_E_*J*_EE_ > *g*_I_*J*_II_ which is harder to satisfy especially when inhibition is fast (*q* is small). Thus, we typically expect that one eigenvalue should decrease (or, at least, its growth should be delayed) before growing again (Figure S2A).

As for the shear force, a similarly simple expression can be obtained in the case of real eigenvalues by noting that the sum of squared eigenvalues in 2 × 2 matrix *𝒥* is equal to (Tr*𝒥*)^2^ — 2Det*𝒥*. This observation yields

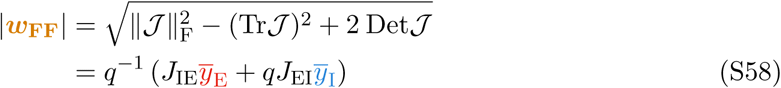

i.e. the shear force is proportional to a weighted average of mean *V*_m_ responses in the E and I units, which, in the SSN, shows linear growth for small *α* and sublinear growth for larger *α* (cf.Section S2 and Figure S1D). Thus, we have a situation in which the force that boosts variability grows faster initially than those that quench variability, causing a transient increase in total variance for small increasing inputs. For large *α*, all forces (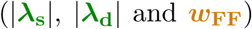) grow as 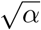 (Figure S2A), because *𝒥* is dominated by its **J**_*αβ*_*y̅_β_* components and the *y̅* terms grow as as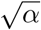 seen in Section S2. Thus, the total variance in Equation (S55) should decay as 1/*α* in this limit, consistent with what we concluded in Section S3.3.

When the eigenvalues of *𝒥* turn complex-conjugate, Equations (S56) to (S58) above become more complicated expressions, which nevertheless does not change the main insights.

## S4 Analysis of the balanced ring network

### S4.1 Reduced Schur decomposition

In this section we describe the mathematical details underlying Figure 5E of the main text. As we did above for the two-population model (Section S3.5), we want to gain some mechanistic understanding of how the input modulates variability in the ring SSN, through an analysis of the “forces” that the network dynamics impose on the flow of fluctuations, thereby affecting noise variability. To study fluctuations, we begin by linearising the dynamics of the network around the fixed point induced by the external input (we fix the motion direction *θ*_s_ to 0° without loss of generality). This leads to the same Equations (S18) and (S19) as above, where the effective connectivity matrix *W*^eff^ (*c*) is now an *N* × *N* matrix that depends on the contrast variable *c* (cf. Equation (8) in the main text). Next, we seek a low-dimensional reduction of those linearized dynamics: we write *δV*(*t*) = Uy(*t*) for some y ∈ ℝ^*K*^ and reduced orthonormal basis U ∈ ℝ^*N*×*K*^ with *K* ≪*N*, and look for dynamics of the form

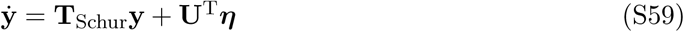

where, for interpretability, **T**_Schur_ ∈ ℝ^*K*×*K*^ is constrained to be quasi-upper-triangular. The covariance matrix **Σ** of δ**V** is then approximated by Σ≈ U cov(y) U^T^, where cov(y) is obtained from standard linear systems theory by solving a reduced-order Lyapunov equation (e.g. Appendix A).

While methods exist that will perform the above model-order reduction to best approximate the covariance of *δ***V**, here we instead want to approximate the high-dimensional flow – i.e. approximate the Jacobian *𝒥*(*c*). A natural way of doing this would be to simply Schur-transform the Jacobian *𝒥*(*c*), and truncate the resulting Schur basis appropriately (e.g. look for the columns of U for which the couplings in **T**_schur_ are non-negligible). Complications arise from the Schur decomposition not being unique: prior to orthogonalizing the eigenvectors of *𝒥*(*c*), we are free to order them in any of *N*! possible ways. This may undermine interpretability, because although there might well exist an ordering that returns a very sparse matrix **T**_Schur_, leading to a parsimonious description of the recurrent dynamics in terms of feedforward interactions between a very small number of modes, we might never find such an ordering (e.g. a random ordering typically leads to a dense matrix **T**_Schur_). Another complication relates to the fact that we would like to “follow” those relevant Schur modes and their interactions as we vary the contrast *c* (cf. Figure 5E, right), so we also require the ordering to lead to interpretable dynamics *across contrast levels.* In some cases, there is a natural choice of ordering, e.g. by decreasing order of the corresponding eigenvalue real parts, that may lead to a very sparse Schur triangle with a nice interpretation (Murphy and Miller, 2009). Here, we found it very challenging to find good exact Schur decompositions by hand, and we instead automatised the process of finding good approximate Schur decompositions, as described below.

Here, we instead adopt the following approach. We capitalise on the fact that bump kinetics capture most of the network fluctuations (cf. fitting procedure in Figure 5A-D; see also the PCA analysis in Figure S4A), so that we expect interactions between these three activity modes to form the dominant part of the recurrent network interactions. Let *N*_E_ = *N*_I_ = *M* (we have one excitatory and one inhibitory neuron at each of *M* = 50 sites on the ring) and let b_1_,b_2_ and b_3_ be the three modes of bump motion (defined in ℝ^*M*^) corresponding to fluctuations in bump location, width, and amplitude, respectively. We first orthonormalize these modes, which leaves b_1_ and b_2_ unaffected but results in slight negative flanks in b_3_ (Figure S4B). We then constrain our truncated Schur basis to be made of 3 pairs of sum-like and difference-like modes of the form:

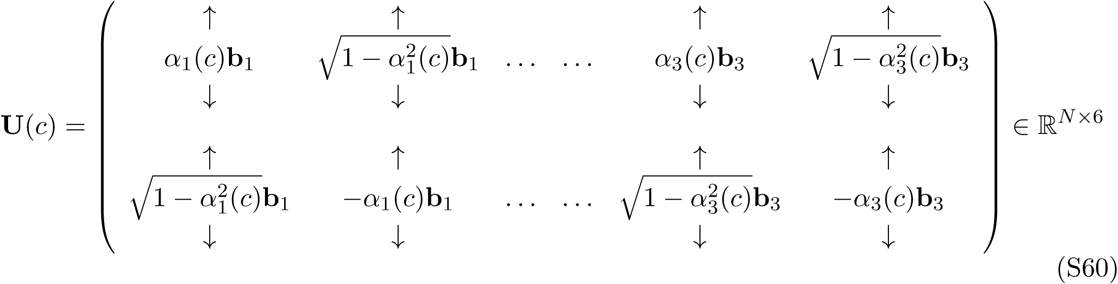

with *α_i_*(*c*) ∈ [0:1] for *i* ∈ {1,2,3}. (Thus, with the notation introduced above, we have *K* = 6 ≪ *N*.) By construction, these modes are orthonormal (**U**^T^**U** = **I** ∈ ℝ^6×6^). We then seek a real quasi-Schur factor with the following structure:

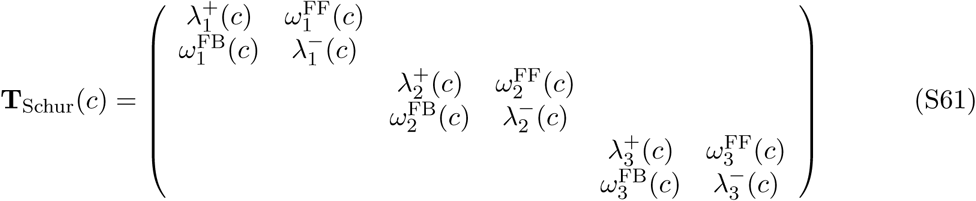

where 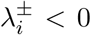 and all non-specified elements are set to zero. We then jointly optimise^8^ both the three *αi* parameters, and all the λ and *ω* parameters (15 parameters in total), to minimise 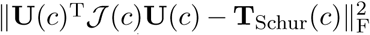. We do this separately for each contrast level, resulting in parameters 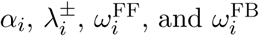 with a smooth dependence on the contrast *c* (Figure S4C).

The green and orange arrows in Figure 5E (left) represent the flow induced by the negative feedback interactions (given by 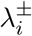 in each plane) and that induced by the feedforward link 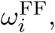 respectively. While we included sum-to-difference feedback terms 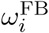 mostly because the real Schur decomposition requires them^9^ and because it seemed to prevent the emergence of degeneracies / local minima in the cost function, we found that they were eventually driven very close to zero. We therefore set them to zero after optimization and in all subsequent analyses. In Figure 5E (middle), green lines show the mean restoring force 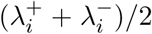 in each plane, while the orange lines show 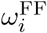 in each plane.

We also checked that the truncation retained the key qualitative aspects of the covariance matrix (Figure S4D). We also tried to fit the full upper-triangular part of **T**_Schur_, but the extra allowed interactions ended up not being used (a single, small feedforward weight from the “amplitude” difference mode to the “width” sum mode was discovered, but it was much smaller than the other feedforward interactions, and setting it to zero did not affect the resulting *V*_m_ covariance matrix qualitatively).

### S4.2 Comparison to a ring attractor model

We compared our ring SSN model to a version of the ring attractor model published in (Ponce-Alvarez et al., 2013). The model was made of a single population with a similar ring topology, and connectivity of the form

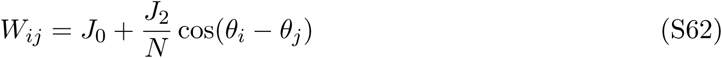

(which violates Dale’s law). The dynamics obeyed a similar Langevin equation as for the ring SSN, namely

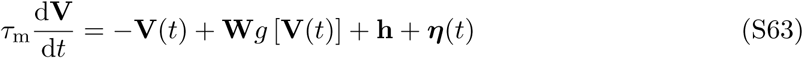

with a saturating firing rate nonlinearity 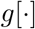 applied pointwise to the elements of **V**,

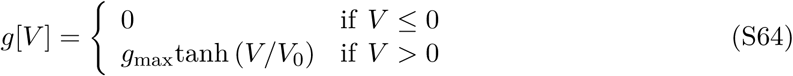

and a noise process *η* identical to the one we used in the SSN (same spatial and temporal correlations), with a variance adjusted so as to obtain Fano factors of about 1.5 during spontaneous activity (Figure S6B, black). The external input had both a DC and a contrast-dependent modulated component: 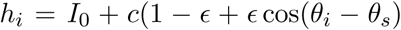 where *θ*_*s*_ is the stimulus direction and *∊* controls the depth of the modulation.

We used the following parameters: *g_max_* = 100, *J*o = −40/gmax, *J*_2_ = 33/gmax, *I*_o_ = 2, *ɛ* = 0.1, and *V*_0_ = 10. Note that although the phenomenology and dynamical regime of this model was consistent with that of Ponce-Alvarez et al. (2013) (Figure S6), the model differed in some of the details: our dynamics were written in voltage form, not in rate form, we have only one unit at each location on the ring (as opposed to small pools), and our input noise process has spatial correlations to allow for a more direct and consistent comparison with the ring SSN.

Our analysis of variability in this ring attractor network is presented in Figure S6 in a format identical to that of Figure 5 of the main text, and shows that shared variability is entirely dominated by the fluctuations in the location of an otherwise very stable bump of activity.

## Appendices

### A Derivation of the total variance in the 2-population model

In this section we derive the result of Equation (S23). We use the fact that the stationary covariance matrix of a process governed by linear stochastic dynamics is given in algebraic form by a Lyapunov equation. Specifically, when the spatial and temporal correlations in the noise term *η* in Equation (S18) are separable, we can augment the state space with two noise units and write their (linear) Langevin dynamics as

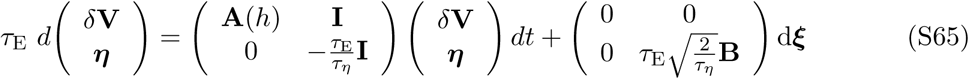

where d*ξ* is a unit-variance, spherical **W**iener process, and **B** is the Cholesky factor of the desired noise covariance matrix, that is, Σ*_η_* = **BB**^T^ (the 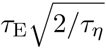 factor is such that this equality holds). Then, from multivariate Ornstein-Uhlenbeck process theory (Gardiner, 1985), we know that the covariance matrix of the compound process satisfies the following Lyapunov equation:

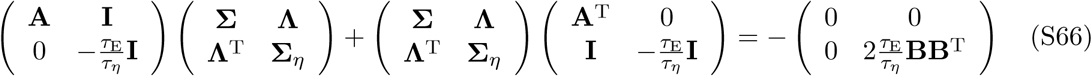

where Σ is the covariance we are trying to compute. By vectorizing Equation (S66), neglecting the bottom right quadrant (which by itself only confirms **Σ_η_** = **BB**^T^ as promised above), and taking into account the symmetry, one ends up with a system of 7 coupled but *linear* equations to solve for the 3 unknowns of **Σ** and the 4 unknowns of **Λ**. This can be done by hand using some patience, or automatically using a symbolic solver such as Mathematica, and yields the expression in Equation (S23).

## Footnotes

1 Talking about how *y̅*E scales with large α actually stops making sense whenΩ_E_<0 precisely because for large enough α the E unit stops firing; but the point here is that because *Y̅*_e_must decrease at some point, it will necessarily become strongly sublinear in α over some range before it starts to decrease;

2 The other limit (fast noise, *τ*_η_ → 0) also greatly simplifies Equation (S27), but would not make much sensein the context of this study, since Equation (S1) is meant to model the dynamics of the voltage on a timescale ≥ 30 ms, which is the timescale on which a threshold power-law relationship between voltage and rate has beenmeasured in cat V1. Therefore, the input noise that we explicitly model here is meant to capture the slowly fluctuating components of external inputs, the fast components having been “absorbed” into the threshold powerlawgain function

3 In this regime, *𝒥*_EE_*y̅*_E_ ξ ⇔1 *A*_EE_ > 0 implies instability of the excitatory subnetwork in isolation, and therefore the need for dynamic, stabilizing feedback inhibition (hence the name “inhibition-stabilized network”).

4 The variance of the effective noise process is proportional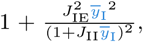 and so has some dependence on α especially for small α before *y̅*_I_ grows large. However, empirically, the quality of the approximation in Equation (S39) – which is derived under the assumption of constant effective noise variance – suggests we can neglect this e

5 The eigenvalues remain real over the entire input range for about half of the 1000 random networks studied throughout (all with *q* = 1/2). In the second half, they go from real to complex-conjugate and then sometimes to real again.

6 This holds when the eigenvalues of A are real. When they are complex conjugate, one can still perform a real Schur decomposition composition by orthogonalizing the imaginary part of the eigenvector against the real part, which yields 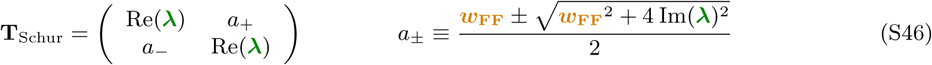 and the two Schur vectors in this case are also sum-like and difference-like in this order. At this point (anticipating a little bit on what follows this footnote), we note that in the imaginary case, there is a small feedback term proportional to *a*_, from the sum-mode to the dfference-mode. Thus, the picture of the forces drawn in Figure 2 of the main text is incomplete. However, we will see that in the slow-noise limit(which gives a very good approximation to the output covariance as seen in Section S3.3), the purely feedforward picture remains exact provided one replaces *ω*_FF_,λ_d_ and λ_r_ by their moduli.

7 More generally, for arbitrary *q*, *κ* and *ρ*_EI_, in the limit *r* → œ, Equation (S55) still holds, in precisely the same form, but in terms of the eigenvalues and feedforward Schur weight of 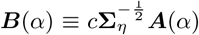 rather than of *𝒥*(*α*). This is because, in that limit, 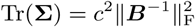. Note that *q* cannot affect the result in the limit *τ*_*η*_ → œ; and that when *κ* = *q* and *ρ_EI_ =* 0, then *𝒥 (*α*) = *B*(*α*)* and hence Equation (S55) holds. To see why 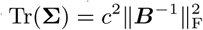 in this limit: most simply, in the slow noise limit, one can think of the noise *η*(*t*) Equation (S18) as a constant input and solve for its steady state 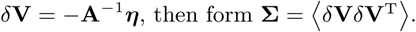

8 We use straightforward re-parameterisation to enforce the constraints 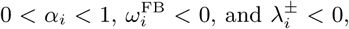 to turn the problem into an unconstrained minimization problem that converges within a few tens of quasi-Newton iterations (BFGS algorithm).

9 When some eigenvalues of J (c) are complex-conjugate, the real Schur decomposition cannot yield an exactly triangular matrix **T**_Schur,_ which will have some 2 × 2 square matrices along its diagonal. Cf. footnote 6 above.

---

**Figure S1:**
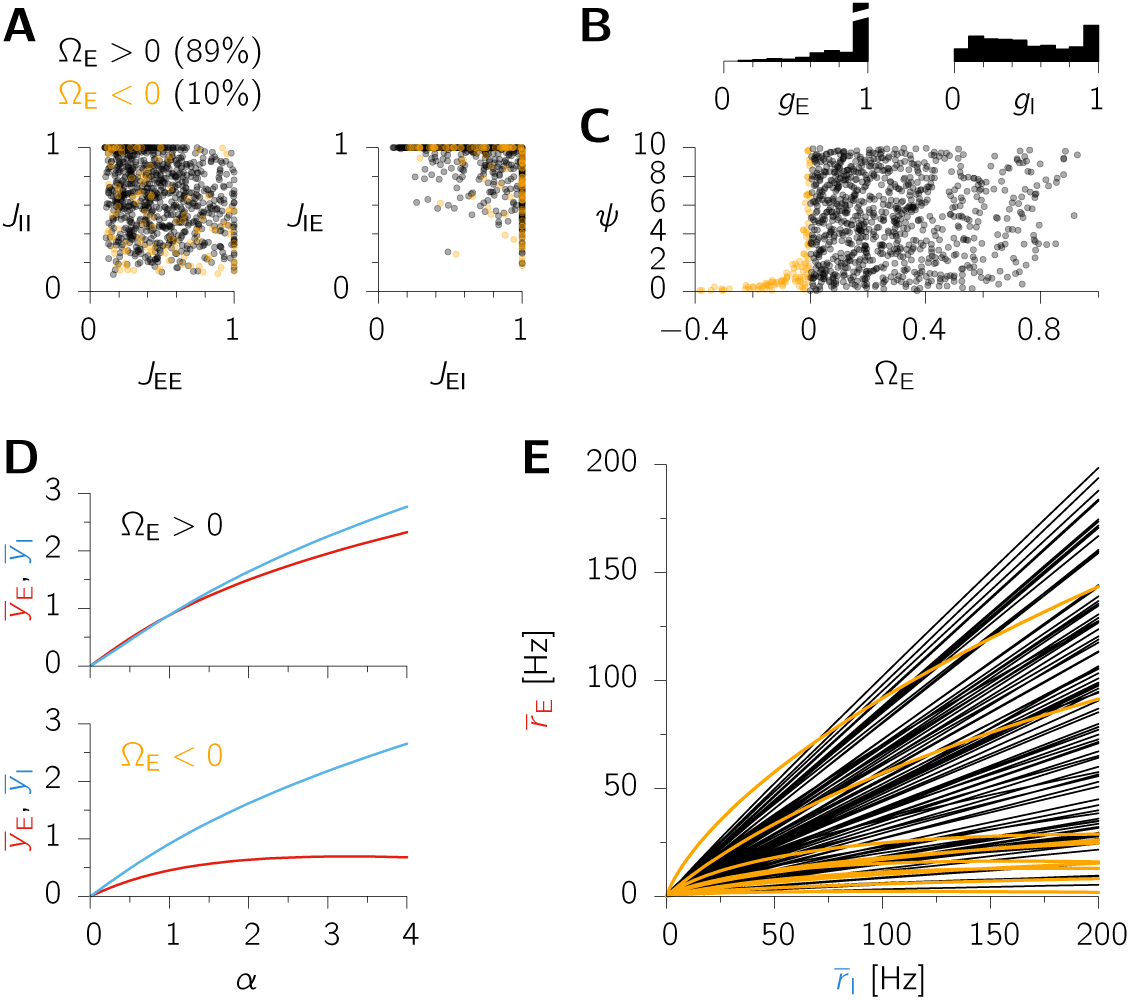
(related to Figure 1 of the main text) – Typical behavior of mean responses to increasing inputs in the 2-population SSN. (**A**) Dimensionless recurrent weights {*J*_*αβ*_} for our 1000 randomly sampled networks; these are normalized such that the largest of the four weights is one. Colors indicate the sign of Ω_E_. (**B**) Distribution of feedforward weights *g*_E_ and *g*_i_, also normalized for each network so that their maximum be one. (**C**) Overall connection strength *ψ* (such that *W_αβ_* ≡ *ΨJ*_αβ_) vs. Ω_E_. (**D**) Example responses (dimensionless voltages *y̅*_E_ and *y̅_I_*) to increasing inputs (dimensionless *α*), for a network with Ω_E_ > 0 (top) and one with Ω_E_ < 0 showing supersaturation (bottom). (**E**) Mean E firing rate *r̅*_E_ as a function of the mean I firing rate *r̅*_I_, for a subset of networks; each point on these curves corresponds to a different input level, increased from zero to a maximum value chosen such that *r̅*_i_ = 200 Hz.

**Figure S2:**
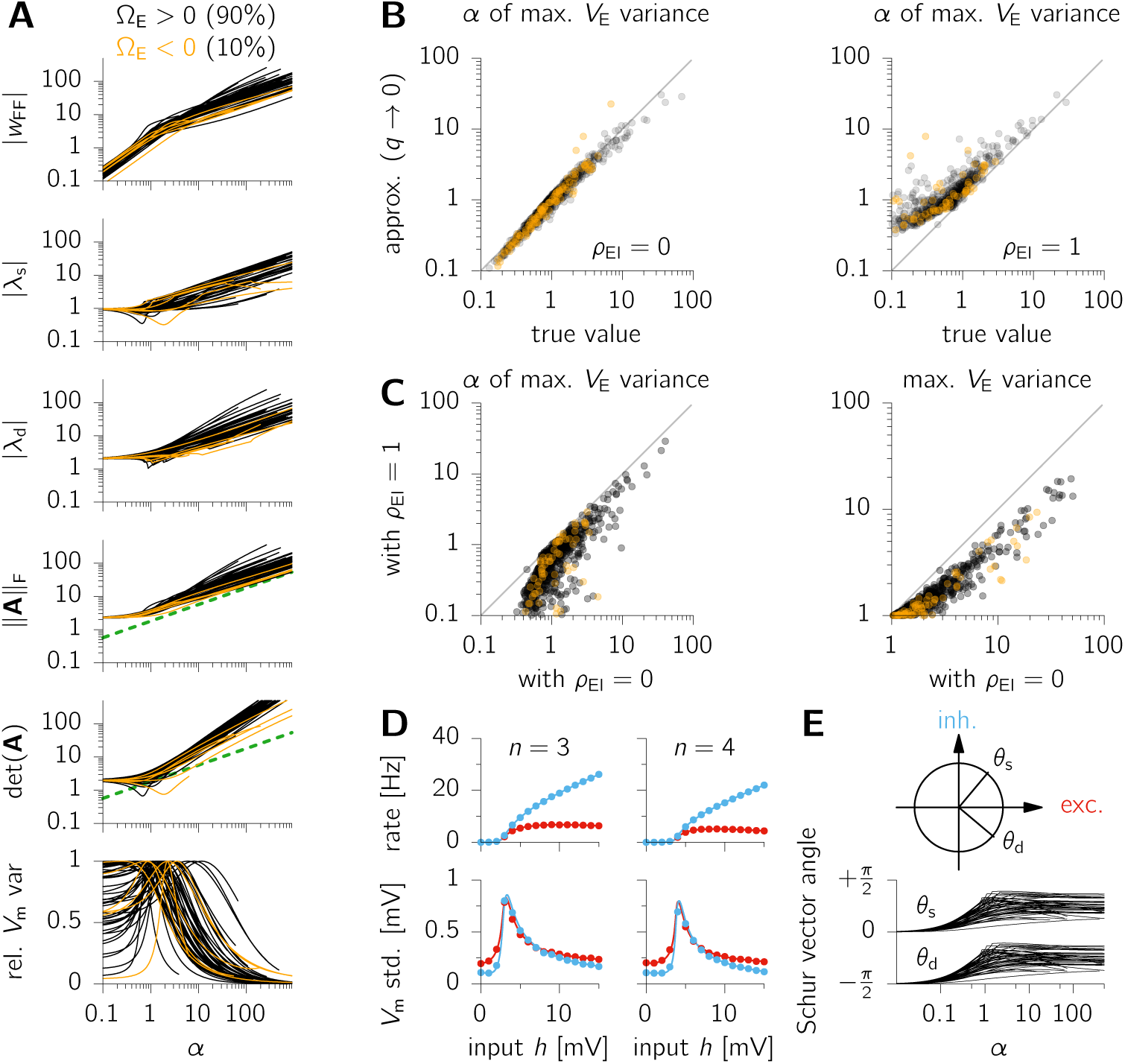
(related to Figure 2 of the main text) - Robustness of variability modulation to changes in network parameters. We examined the modulation of variability by external input in the 1000 randomly parameterized, 2-population networks of Figure S1.(**A**) Behavior of 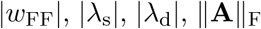, det(A) and the total variance (normalized to unit peak), as a function of the (dimensionless) input *α*. The dashed green line is proportional to 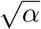 Only a random subset of the thousand random networks are shown. Following the same convention as in Figure S1, cases with Ω_E_ > 0 are shown in black, those with Ω_E_ < 0 in orange. (**B**) Scatter plot of the *α* at which the E variance reaches its maximum (“true value”), and that given by the approximate criterion of Equation (S40) (which assumes very fast inhibition, i.e. *q* − 0), for uncorrelated (left, *ρ*_EI_ = 0) and fully correlated (right, *ρ*_EI_ = 1) input noise term to the E and I units.(**C**) Scatter plot of the input *α* at which the E variance peaks (left), as well as the value of the variance peak (right), for *ρ*_EI_ = 0 vs. *ρ*_EI_ = 1. (D) Mean E (red) and I (blue) firing rates (top) and *V*_m_ std. (bottom) for larger values of the power-law exponent *n*; parameters were otherwise the same as in Figure 1 of the main text. (**E**) Orientation of the two Schur vectors for a subset of the 1000 random networks. Their “sum-like” and “difference-like” nature emerges quite rapidly for small *α* and then persists for larger *α*.

**Figure S3:**
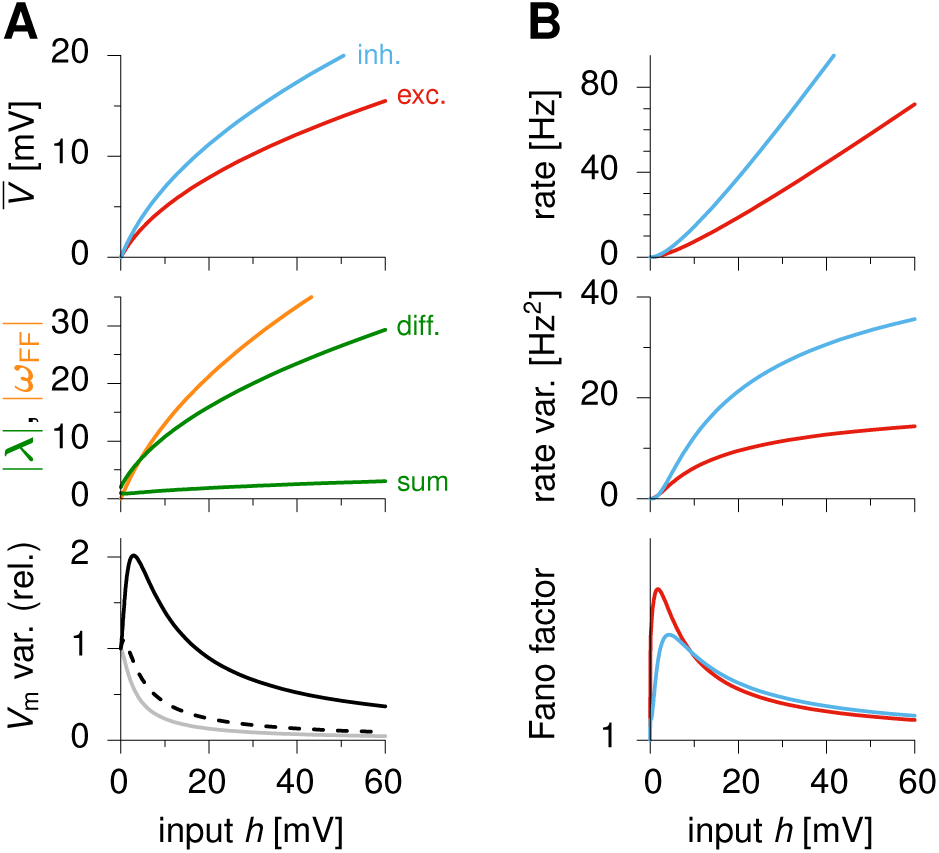
(**A**) Example network showing transient increase in variability with increasing external input *h* (black), *without* any substantial decrease in |λ_s_| (lower green). The dashed black line shows the predicted variability (Equation (S55)) assuming *ω*_FF_ = 0 uniformly, i.e. taking into account only the magnitude of the restoring forces λ_d_ and λ_s_. The gray line is the prediction made by assuming fully correlated input noise terms with variance 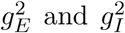 respectively for the E and I units. Variability in this case can be read off the slope of the *V*_E_ and *V*_j_ curves (top), because input noise becomes equivalent to fluctuations in *h* to which the network has time to respond. Neither of these two cases correctly predict the initial growth of variability. (**B**) Mean firing rates (top), variances of firing rate fluctuations (middle) and Fano factor (assuming Poisson spike emission on top of rate fluctuations), in the same network as in (A). Note that the overall scale of super-Poisson variability (Fano factor minus one) is arbitrary here, and in general depends on the counting window, autocorrelation time constants, and the variance of the input noise. Parameters: *τ_η_*−,*g*_E_ = 0.77,*g*_I_ = 1,*J*_EE_ = 0.38,*J*_ej_ = 0.27,*J*_IE_ = 1, *J*_II_ = 0.6, *ψ* = 2.37.

**Figure S4:**
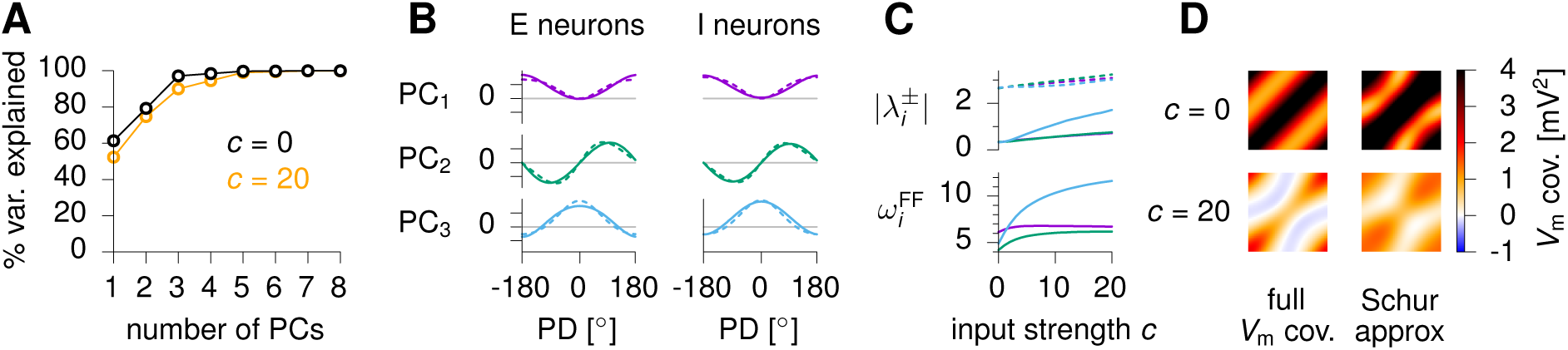
Approximating balanced ring dynamics in a low-dimensional subspace.(**A**) PCA analysis on spontaneous *V*_m_ activity (*c* = 0, black) and high-contrast evoked activity (*c* = 20, orange); shown here is the cumulative percentage of variance explained by an increasing number of retained principal components. Our of 100 components, between 3 and 5 components are enough to explain more than 90% of the *V*_m_ variance. (**B**) The top three PCs in evoked activity (*c* = 20; solid lines) are almost identical to the 3 (orthonormalized) modes of joint E/I bump kinetics (dashed lines). (**C**) Contrast dependence of the 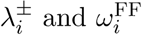 parameters, as obtained from the optimization procedure described in the text, which aims at finding the most accurate, low-dimensional approximation to the (contrast-dependent) Jacobians in Schur form.(**D**) Full *V*_m_ covariance matrix Σ (left) compared with the covariance matrix Σ˜ obtained from the low-dimensional projection of the network dynamics as explained in the text (right), for *c* = 0 (top) and *c* = 20 (bottom). Only the excitatory-excitatory part of the covariance matrices are represented here, but the other 3 quadrants are equally well approximated.

**Figure S5:**
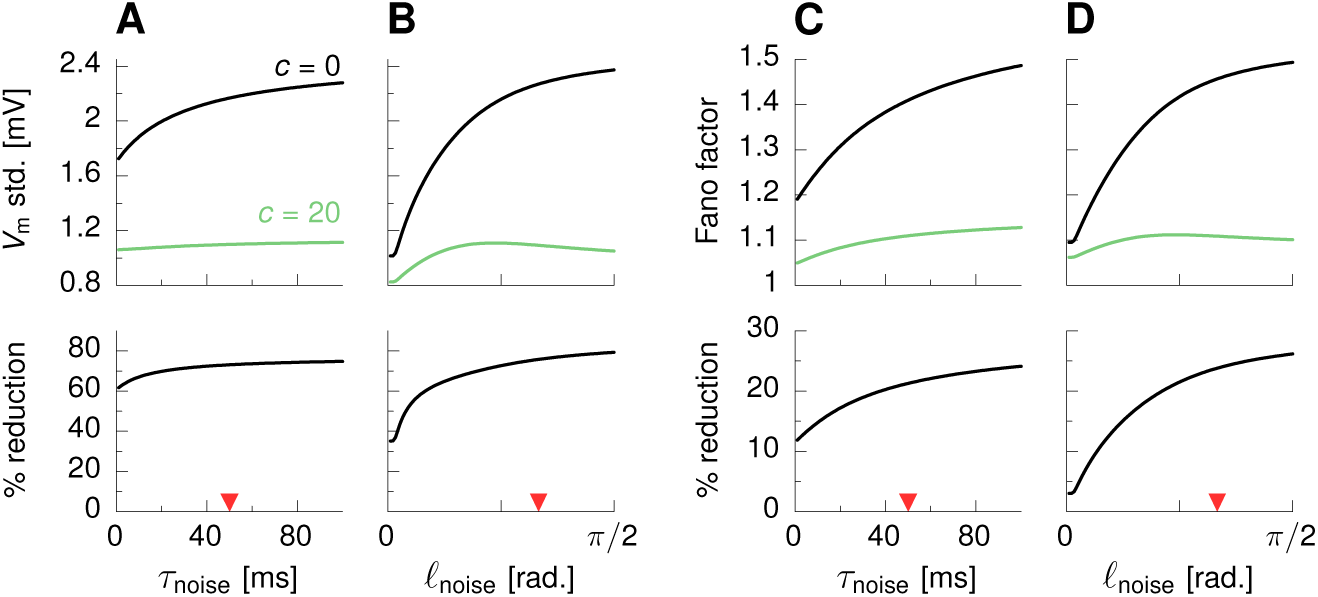
Variability reduction in the ring SSN model depends on spatial and temporal correlations in the input noise. Dependence of the network-averaged *V*_m_ std. (A-B) and Fano factor (C-D) on either the temporal correlation time constant *τ*_noise_ in the external input noise term (for fixed 𝓁_noise_ = 60°), or its spatial correlation length 𝓁_noise_ (for fixed *𝓁*_noise_ = 50 ms), in the spontaneous (*c* = 0, black) and high-contrast (*c* = 20, green) input regimes. Red arrows indicate the nominal parameter values used in the main text. The bottom row shows the amount of relative variability suppression, as a percentage of the mean spontaneous variability.

**Figure S6:**
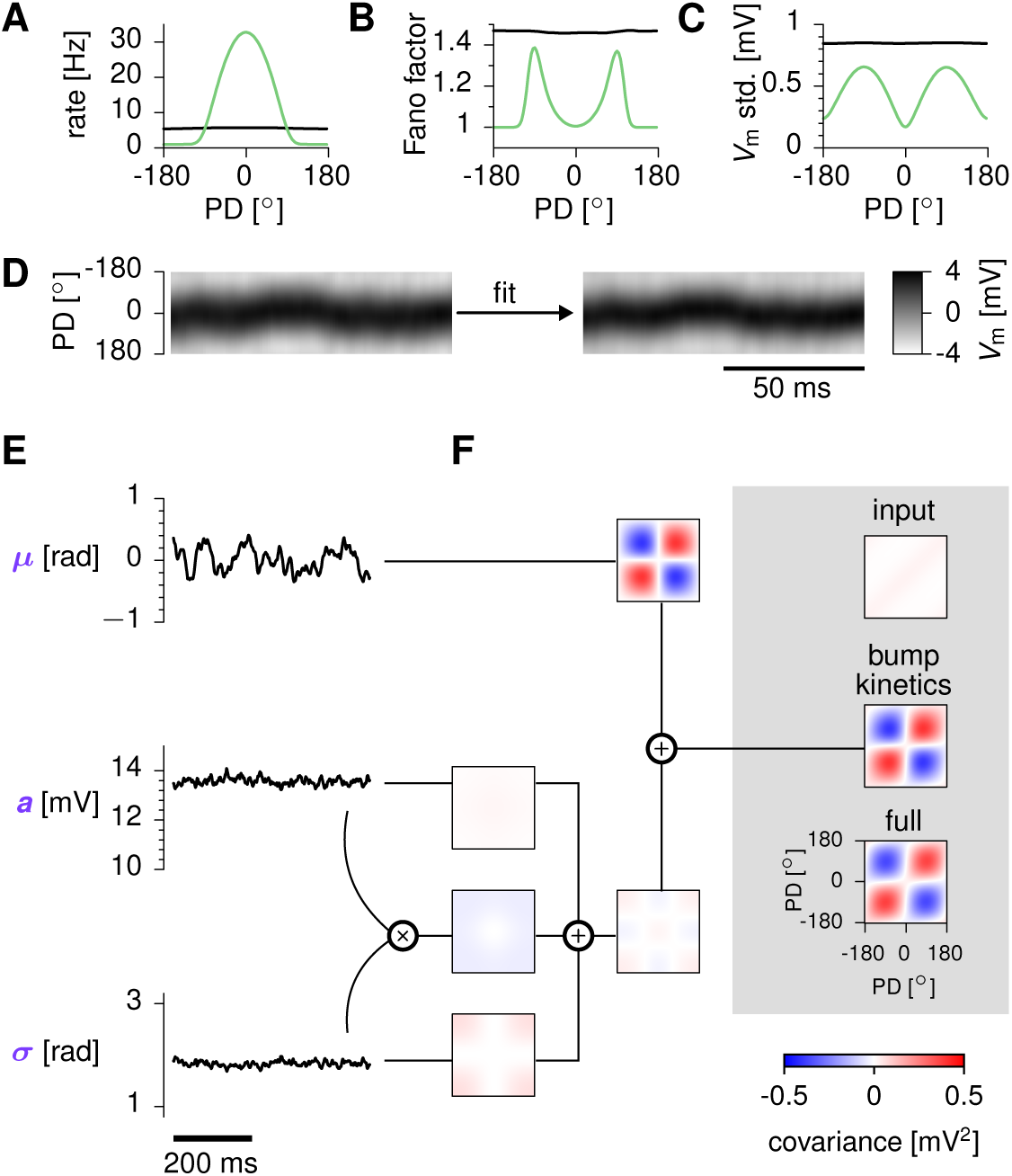
Activity variability in a ring *attractor* network (related to Figure 5 of the main text). (**A**–**C**) Tuning of mean firing rates, Fano factors, and *V*_m_ std. in spontaneous (*c* = 0, black) and evoked (*c* = 3, green) conditions. (**D**–**F**) Analogous to Figure 5D-F of the main text, for the ring *attractor* network. The main contributor to activity variability in this attractor network for strong stimulus is the sideways jittering of the activity bump.

